# Shiba: A versatile computational method for systematic identification of differential RNA splicing across platforms

**DOI:** 10.1101/2024.05.30.596331

**Authors:** Naoto Kubota, Liang Chen, Sika Zheng

## Abstract

Alternative pre-mRNA splicing (AS) is a fundamental regulatory process that generates transcript diversity and cell type variation. We developed Shiba, a comprehensive method that integrates transcript assembly, splicing event identification, read counting, and differential splicing analysis across RNA-seq platforms. Shiba excels in capturing annotated and unannotated AS events with superior accuracy, sensitivity, and reproducibility. It addresses the often-overlooked issue of junction read imbalance, significantly reducing false positives to aid target prioritization and downstream analyses. Unlike other tools that require large numbers of biological replicates or resulting in low sensitivity and high false positives, Shiba’s statistics framework is agnostic to sample size, as demonstrated by simulated data and its effective application to real *n*=1 RNA-seq datasets. To extend its utility to single-cell RNA-seq, we developed scShiba, which applies Shiba’s pseudobulk approach to analyze splicing at the cluster level. scShiba successfully revealed AS regulation in developmental dopaminergic neurons and differences between excitatory and inhibitory neurons. Both Shiba and scShiba are available in Docker/Singularity containers and Snakemake pipelines, ensuring reproducibility. With their comprehensive capabilities, Shiba and scShiba enable systematic quantification of alternative splicing events across various platforms, laying a solid foundation for mechanistic exploration of the functional complexity in RNA splicing.

## Introduction

High-throughput RNA sequencing (RNA-seq) has been instrumental in unraveling the complexity of transcriptomes generated by alternative pre-mRNA splicing and processing. Regulatory functional splicing events play a pivotal role in shaping proteome diversity, cell type transitions, and cellular responses (1–3). The accurate detection and quantification of differential alternative splicing events from RNA-seq are fundamental prerequisites for downstream dissection of regulatory mechanisms and mechanistic characterization of functional splicing.

Several computational tools have been developed to analyze alternative splicing events from RNA-seq data, categorized into exon-centric and intron-centric tools. Exon-centric tools, such as rMATS (4, 5), SUPPA2 (6), Whippet (7), MISO (8), Vast-tools (9), JUM (10), and Quantas (11), examine differential splicing patterns at the exon level for changes in exon inclusion or exclusion across biological conditions. Intron-centric tools, such as MAJIQ (12, 13) and LeafCutter (14), delve into cross-intron splice site usages. Intron-centric tools are specialized in detecting complex splicing junctions, which can be overlooked by exon-centric analyses. On the other hand, exon-centric analyses derive intuitive differential splicing values (ΔPSI), which offer a tangible path to interpret the functional outcomes of alternative splicing regulation. The choice between exon-centric and intron-centric tools depends on specific research questions and the complexity of the alternative splicing events under investigation.

Despite their utility, results from these existing tools often have limited overlap (15), reflecting the diverse statistical approaches employed by each method. This partial intersection of analysis results highlights the different strengths and focuses of each tool, allowing researchers to choose the most appropriate method for their specific needs in the study of alternative splicing events. For researchers outside the RNA biology field, particularly those focusing on the functional aspects of alternative splicing regulation, high-confidence results are most crucial for guiding downstream analyses. Most existing pipelines apply statistical methods requiring many biological replicates, sometimes significantly higher than experimental practicality, to claim differentially spliced events (DSEs), otherwise resulting in relatively low sensitivity and high false positives. Many public bulk RNA-seq data with limited or lacking replicates are therefore excluded from rigorous splicing analyses to derive high-confidence information. Second, existing tools neglect junction read imbalance, leading to a further increase in false positives and inaccurate interpretation of splicing events. Additionally, statistical approaches of existing pipelines are not applicable to single-cell/nucleus RNA-seq (sc/snRNA-seq) data. Finally, some existing tools, particularly those following an exon-centric approach, rely predominantly on annotated splicing events. This limitation results in the oversight of unannotated and cryptic splicing events.

The increasing amounts of sc/snRNA-seq data present a gap in developing a useful statistical framework to extract DSEs as reliably as possible. Several tools, such as Expedition (16), SCATS (17), DESJ-detection (18), BRIE (19, 20), Psix (21), and MARVEL (22), have been developed to explore splicing variability across individual cells at a single-cell level. However, depending on read coverage per cell, their practical uses are limited by current sc/snRNA-seq technologies. Popular scRNA-seq platforms, such as the droplet-based 10x technology, exhibit severely biased and non-uniform read coverage as well as substantial dropouts.

Despite the limitation in read coverage, excluding scRNA-seq data from splicing analysis would be a missed opportunity, especially given the growing volume of data. An alternative use of sc/snRNA-seq data for extracting splicing information would be aggregating reads at the cell cluster level to increase sequencing depth. This approach directly addresses the low coverage issue but presents a *n*=1 problem for statistical analyses of differential splicing, because each cell cluster is defined only once.

In this study, we present Shiba, a novel computational method specifically designed to fill the gaps in current splicing analysis tools and enhance the precision of calling differential alternative splicing using either bulk or sc/snRNA-seq data. Shiba incorporates advanced methodologies for reference-guided transcript assembly, accurate identification of both annotated and unannotated splicing events, and a meticulous statistical framework for differential splicing analysis. By leveraging these improvements, Shiba provides researchers a comprehensive tool with high confidence for unraveling the complexity of alternative splicing dynamics in diverse biological contexts. Applicable to both bulk RNA-seq and scRNA-seq, Shiba is a versatile approach for cross-platform analyses. Through rigorous performance evaluations and comparisons with established tools, we herein demonstrate the superior efficiency and accuracy of Shiba as a valuable tool for the investigation of RNA splicing regulation.

## Materials and methods

### Method overview

Shiba comprises four steps: transcript assembly, detection of alternative splicing events, extraction of splice junction reads, and identification of differentially spliced events (DSEs). During the first step, mapped RNA-seq reads are assembled to build transcript structures using StringTie2 (version 2.2.1) (23). The “-G reference.annotation.gtf --conservative” option is used for each file, and the transcripts are merged with the “-G reference.annotation.gtf --merge” option. This step helps discover unannotated splicing events not described in the reference gene annotation file. In the second step, alternative splicing events are extracted from the newly assembled transcript structures and classified into eight classical splicing events: skipped exon (SE), alternative five prime splice site (FIVE), alternative three prime splice site (THREE), mutually exclusive exons (MXE), retained intron (RI), multiple skipped exons (MSE), alternative first exon (AFE), and alternative last exon (ALE). In the third step, the per-sample number of exon-exon and exon-intron junction reads are counted using RegTools (version 1.0.0) (24) and featureCounts implemented in the SubRead package (version 2.0.3) (25), respectively. In the fourth step, junction reads of replicated samples for each group are pooled together, and two-by-two tables of the count of inclusion and exclusion reads for target exons in two groups are created. Two two-by-two tables are made for SE and RI for both sides of exon-exon/exon-intron junctions. One table is made for FIVE, THREE, AFE, and ALE. Four tables are made for MXE. *n*+1 tables are made for MSE with *n* skipped exons. For a given SE or RI event with enough read coverage (Either >20 on inclusion or >10 on exclusion junction in both groups by default), odds ratio (OR) and Percent Spliced In (PSI) are calculated as follows:

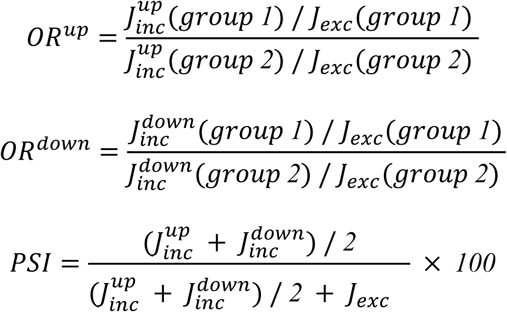

where 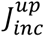 and 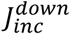 are the number of inclusion junction reads for upstream and downstream, respectively, and *J*_*exc*_ is the number of exclusion junction reads.

For a given FIVE, THREE, AFE, or ALE event with enough read coverage, OR and PSI are calculated as follows:

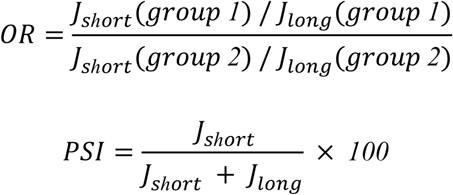

Where *J*_short_ is the number of junction reads spanning the shorter intron, and *J*_*long*_ is the number of junction reads spanning the longer intron.

For a given MXE event with sufficient read coverage, OR and PSI are calculated as follows:

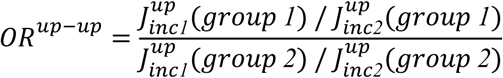

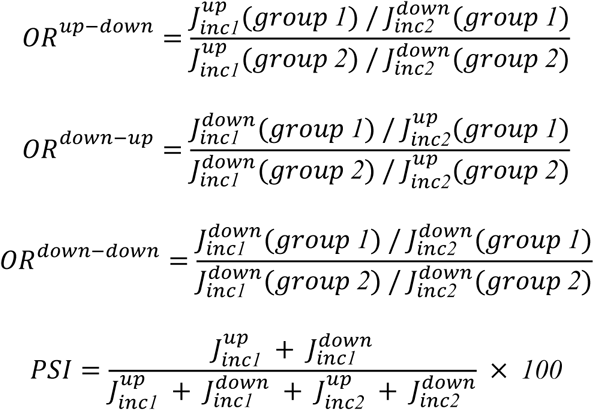

where 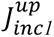 and 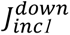 are the number of upstream/downstream inclusion junction reads for the upstream exon, and 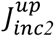 and 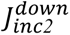 are the number of upstream/downstream inclusion junction reads for the downstream exon.

For a given MSE event with *n* skipped exons, the *OR*^*k*^ for each inclusion junction *k* is calculated as follows:

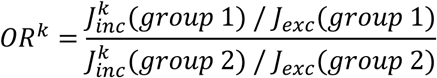

where 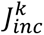 and *J*_*exc*_ are the number of inclusion junction *k* and exclusion junction reads. The index *k* represents each inclusion junction in the set:

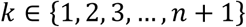

PSI is calculated as follows:

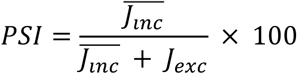

where 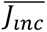 is the average inclusion junction read count, calculated as:

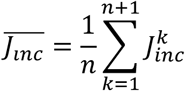

Fisher’s exact test is performed for each two-by-two table. The *P*-values were adjusted based on the false discovery rate using the Benjamini–Hochberg method (26). DSEs are defined as those that satisfy all of the following criteria:

- All OR > 3/2 or All OR < 2/3
- Adjusted Fisher’s exact test *P* (*q*) < 0.05
- |ΔPSI| > 10

Users have the flexibility to configure Shiba parameters, such as the minimum read coverage, *q*-value cutoff, and ΔPSI, according to their preferences. In cases with multiple replicates for two groups, Shiba conducts Welch’s t-test for PSI values (referred to as Shiba+), providing a comprehensive assessment to identify DSEs across multiple metrics. In addition to splicing analysis, Shiba incorporates standard gene expression analysis by quantifying mapped reads per gene using featureCounts. This is followed by normalization and differential expression analysis using DESeq2 (version 1.34.0) (27) and simultaneous calculation of TPM and CPM values, offering additional metrics for diverse downstream analyses.

Shiba has an alternative version of splicing analysis using sc/snRNA-seq data, named scShiba. The processing steps are similar to those of Shiba: Detection of alternative splicing events, pooling splice junction reads for each cell type, and identification of DSEs. First, alternative splicing events are extracted from the gene annotation file and classified into seven classical alternative splicing events (SE, FIVE, THREE, MXE, MSE, AFE, and ALE). Second, taking the junction read count matrix produced by STARsolo (28) and a table of cell barcodes and the corresponding cell types as inputs, reads mapped to each junction are pooled for each cell type so each cell type can be treated as a biological replicate in the following statistical analysis. Finally, the PSI of each splicing event is calculated, and differential splicing analysis is performed in the same way as in Shiba.

All the necessary software and scripts required to run the Shiba/scShiba pipeline are provided as a single Docker/Singularity container, ensuring a highly reproducible analysis environment. Furthermore, we offer the SnakeShiba pipeline, based on the Snakemake workflow (29), facilitating accelerated execution through optimized parallelization.

### Simulated RNA-seq data

We executed SnakePolyester (version 0.1.0) (https://github.com/NaotoKubota/SnakePolyester), a Snakemake workflow for simulating RNA-seq reads by Polyester (version 1.22.0) (30), to generate control and perturbed RNA-seq samples (each with 101 bp paired-end reads). Different base read coverages (low, moderate, high, and very high), biological replicates (ranging from 1 to 10), and variations between replicates (low, moderate, and high) were set. The parameters in a configure file were as follows: For base read coverage, “coverage” is 10 (low), 20 (moderate), 30 (high), 100 (very high). For variation between replicates, “size_factor” is 0.3 (low), 10 (moderate), and 30 (high). For performance evaluation, we randomly selected 3,000 mouse protein-coding genes with alternative splicing events for SE (1,750), FIVE (500), THREE (500), MXE (50), and RI (200). For AFE and ALE evaluation, we selected 2,400 genes (1,200 each), and for MSE, another 2,400 genes with MSE events. We divided the genes into three groups: (i) genes with unaltered splicing, (ii) genes with 20% PSI increase of an alternative splicing event, and (iii) genes with 20% PSI decrease of an alternative splicing event in the perturbed samples. These groups were used as the ground truth to evaluate the performance of various computational tools for alternative splicing analysis. The simulated reads were generated in FASTA format and converted to FASTQ format using seqtk seq function (version 1.4) (https://github.com/lh3/seqtk) with the “-F I” options. These reads were mapped to the mouse reference genome (mm10) and the Ensembl 102 transcript annotations using STAR (version 2.7.10b) (31) with the “--outFilterMultimapNmax 1” option. To evaluate the Shiba performance at detection of unannotated alternative splicing events, we customized the reference gene annotation file by removing records of inclusion isoforms.

### Real RNA-seq data

For the performance evaluation, RNA-seq samples from mouse adult liver and kidney tissues (32) were downloaded from GEO (GSE90179 and GSE90180). Each tissue has six replicates, divided into two biological replicates that consist of three technical replicates. We performed quality control using fastp (version 0.23.2) (33) with the “-3 -q 30” options to discard low-quality reads. We aligned the remaining reads to the mouse reference genome (mm10) and the Ensembl 102 transcript annotations using STAR with the “--outFilterMultimapNmax 1” option.

For the evaluation of run time and junction read imbalance, RNA-seq samples from cytoplasmic fraction of mouse embryonic stem cells (mESC) with siRNA-mediated double knockdown (KD) of *Ptbp1* and *Ptbp2* were downloaded from GEO (GSE159993) (34). We performed quality control using fastp (33) and aligned the remaining reads to the mouse reference genome (mm10) using STAR with the same options.

For the comparison of RNA splicing quantification between RNA-seq and RT-PCR, RNA-seq samples of two prostate cancer cell lines, PC3E and GS689 (4) (SRP014759), and a cultured T-cell line, Jurkat, with KD of *hnRNPL* (35) (SRP059357) were downloaded from SRA. The reads were aligned to the human reference genome (hg19) Ensembl 87 transcript annotations using STAR with the “--outFilterMultimapNmax 1” option after quality control by fastp. These datasets were then used to evaluate the accuracy of exon-level PSI quantification. The PSI values for the tested skipping exons were obtained from the corresponding studies, and Pearson’s correlation coefficient (r) was calculated to compare the estimates between RT-PCR and RNA-seq data.

For the identification of alternative exons associated with nonsense-mediated mRNA decay, RNA-seq samples of HEK293 Flp-In T-Rex cells with siRNA-mediated double KD of *XRN1* and *UPF1* were downloaded from GEO (GSE57433) (36). We aligned the high-quality reads to the human reference genome (hg38) and Ensembl 106 transcript annotations using STAR with the “--outFilterMultimapNmax 1” option. The mapped reads were analyzed by Shiba with the default settings to identify DSEs between the control and the KD sample.

For the identification of neuronal cell class-specific alternative splicing events, RNA-seq of tagged-ribosomal affinity purification (RiboTRAP) samples from genetically-defined neurons were downloaded from GEO (GSE133291) (37). The RiboTRAP samples were generated using conditionally HA-tagged endogenous ribosomal protein Rpl22 in glutamatergic (excitatory) neurons (CamK2-cre and Scnn1a-cre), and GABAergic (inhibitory) interneurons (SST-cre, PV-cre and VIP-cre) in cortex. Within the hippocampus, excitatory neurons (CamK2-cre and Grik4-cre) and inhibitory interneurons (SST-cre) were targeted. Quality control and mapping to the mouse reference genome (mm10) were performed using fastp and STAR with the same option as the other dataset. The mapped reads were analyzed by Shiba with the default settings to calculate PSI for each sample.

For the identification of alternative splicing events under the perturbation of neuronal RNA binding proteins (RBPs), we used RNA-seq of Emx1-cre *Ptbp2* knockout (KO) mouse cortex (postnatal day 1) (GSE84803) (38), Nestin-cre *Mbnl1* and *Mbnl2* double KO mouse adult cortex (SRP142522) (39), and *Mbnl2* KO mouse adult hippocampus (2–3 months old) (SRP013558) (40). Quality control and mapping to the mouse reference genome (mm10) were performed using fastp and STAR with the same option as the other dataset. The mapped reads were analyzed by Shiba with the default settings to identify DSEs between the control and the RBP knockout samples. Enrichment analysis of RBPs’ target events and DSEs during neuronal maturation was performed based on Fisher’s exact test and the *P*-values were adjusted based on the false discovery rate using the Benjamini–Hochberg method.

### Single-cell RNA-seq data

We downloaded the single-cell RNA-seq (scRNA-seq) data of the mouse midbrain dopaminergic neurons (embryonic day 13.5, 15.5, 18.5, postnatal day 1, 7, and 90) from GEO (GSE116138) (41). The raw sequencing reads were aligned to the mouse reference genome (mm10) using STAR in STARsolo mode (28) to obtain the gene expression matrix (GEM) across genes and individual cells and the splice junction matrix (SJM) across junctions and cells. The GEM was processed by Scanpy (version 1.9.5) (42). Possible outlier cells expressing non-neuronal gene markers were removed, and 987 cells were kept for downstream analysis. The table of cell barcodes was used for subsequent scShiba analysis with the SJM to identify DSEs between time points.

### Single-nucleus RNA-seq data

We downloaded the single-nucleus RNA-seq (snRNA-seq) data of the mouse primary visual cortex (postnatal day 28) from GEO (GSE190940) (43). The raw sequencing reads were aligned to the mouse reference genome (mm10) using STAR with the STARsolo mode (28) to obtain the GEM across genes and single nuclei and the SJM across junctions and nuclei. GEM was processed by Scanpy (version 1.9.5) (42). We used Scrublet (44) for cell doublets detection and Harmony (45) for batch-effect correction. Cell types were manually assigned based on the expression level of marker genes. The table of cell barcodes and assigned cell types was used for the subsequent scShiba analysis with the SJM to identify DSEs between cell types of interest.

### HITS-CLIP data

To map RBP-RNA interactions, we downloaded raw HITS-CLIP data of PTBP2 from mouse cortex (embryonic day 18.5) (GSE103315) (46), MBNL1 for mouse whole brain (16 weeks old) (GSE39911) (47), and MBNL2 from mouse hippocampus (2–3 month old) (GSE38497) (40). These data were processed by the Snakemake workflow “preprocessing_HITSCLIP.smk” in SnakeNgs (version 0.2.0). In brief, the raw reads were trimmed by fastp (version 0.23.4) with the “-l 20 -3 --trim_front1 5” options, followed by alignment of the remaining reads to the mouse reference genome (mm10) using STAR (version 2.7.11a) with the “--outFilterMultimapNmax 100” option. Deduplication of mapped reads was performed by Picard (version 3.1.1) (https://broadinstitute.github.io/picard/).

### Performance evaluation using simulated RNA-seq data

RNA-seq reads were analyzed by Shiba (version 0.4.1), rMATS (version 4.3.0) (4, 5), SUPPA2 (version 2.3) (6), Whippet (version 1.6.1) (7), MAJIQ (version 2.4.dev102+g2cae1507) (12, 13), and LeafCutter (version 0.2.9) (14) to call DSEs. Tools except for Shiba were executed by Snakemake workflows in SnakeNgs (version 0.2.0) (rMATS.smk, SUPPA2_diffSplice.smk, Whippet.smk, MAJIQ.smk, and LeafCutter.smk). We ran tools supporting multi-core usage on 16 CPUs, and others on 1 CPU.

#### Shiba

For Shiba, mapped reads and the Ensembl 102 transcript annotation file were used with the default parameters (*q* < 0.05 and |ΔPSI| > 10). Minimum read coverage for differential splicing analysis was set to 10. For Shiba+, in addition to Shiba’s criteria, events that passed Welch’s t-test *P* < 0.05 were called as DSEs when number of replicates are more than two in both groups.

#### rMATS

For rMATS, mapped reads and the Ensembl 102 transcript annotation file were used with the default parameters (*P* < 0.05 and |ΔPSI| > 10). The options “--novelSS --mil 0 --mel 10000” were set to enable detection of unannotated splicing sites.

#### SUPPA2

Salmon (version 1.10.1) (48) was executed to estimate transcript abundance and SUPPA2 was executed with the default parameters (*P* < 0.05 and |ΔPSI| > 10) using the estimated transcript abundance.

#### Whippet

According to the authors’ guidelines, incorporating bam files when creating an index enables the detection of unannotated splicing in Whippet. However, errors occurred during index creation in multiple samples, making analysis impossible. The authors acknowledged this issue as a significant bug (referenced here: https://github.com/timbitz/Whippet.jl/issues/79). Consequently, we opted for standard index creation and restricted our analysis solely to annotated events in this study. We performed analyses using default parameters (Probability > 0.95 and |ΔPSI| > 10) with the FASTQ file as input. Minimum read coverage for differential splicing analysis was set to 10.

#### MAJIQ

Mapped reads and the Ensembl 102 transcript annotation file were used to build splicegraph by “majiq build” command, followed by “majiq deltapsi” or “majiq heterogen” to perform differential splicing analysis with the default parameters. The output files were summarized by “voila modulize” with the “--show-all” option to get statistics of all tested events in the output file “junctions.tsv”. For “MAJIQ deltapsi”, a posterior probability > 0.95 and |ΔPSI| > 10 were set as DSE criteria. For “MAJIQ het”, Welch’s t-test *P* < 0.05 and |ΔPSI| > 10 were set.

#### LeafCutter

For LeafCutter, leafcutter_ds.R was executed to perform differential splicing analysis with the threshold adjusted *P* < 0.05 and |ΔPSI| > 10. Minimum read coverage was set to 10 and minimal samples per intron/group was set to 1.

To assess how well each tool predicted DSEs from simulated data, we used known sets of the differentially spliced and unaltered genes as the ground truth. Since each tool defines alternative splicing events in unique ways, making direct event-to-event comparisons difficult, we performed a gene-level performance evaluation. A gene was considered positive if it contained at least one DSE. We assessed the performance of each tool using three metrics: the false positive rate (FPR), false negative rate (FNR), and Matthews correlation coefficient (MCC). To evaluate performance in detecting unannotated splicing events, we customized the reference gene annotation file by removing records of inclusion isoforms and used it for each computational tool so that alternative splicing events were treated as unannotated events in each analysis.

### Performance evaluation using real RNA-seq data

The execution time of each tool was measured using the Linux time command (/usr/bin/time). We ran each tool for RNA-seq samples from the cytoplasmic fraction of mESC with siRNA-mediated double KD of *Ptbp1* and *Ptbp2* (GSE159993). The control and KD groups have three replicates with 40-60 million reads each.

To evaluate the reproducibility of each tool, we calculated the reproducibility ratio (RR) metric using RNA-seq samples of mouse adult liver and kidney tissues (32). RR quantifies the proportion of DSEs identified by algorithm A in dataset D that are consistently identified using a similar replicate dataset D’, thus representing the internal consistency of the tool. For our RR analysis, we randomly subsampled either one or three replicates from six available replicates per condition (liver and kidney), resulting in nine instances of DS analysis. We then computed the overlap fraction of the top *n* DSEs ranked by statistical metrics (e.g. *P*-value, probability) across the nine analyses (36 comparisons in total). The results were plotted as average RR curves with 95% confidence intervals.

Since highly biased methods can also exhibit high RR, we further calculated the intra-to-inter ratio (IIR) to assess the type I errors of the tools. IIR is an approximation of false discovery proportion. The IIR is determined by comparing the number of DSEs identified in intra-condition comparisons (liver vs. liver) to those found in inter-condition comparisons (liver vs. kidney). Since DSEs identified in intra-condition comparisons are likely false positives, lower IIR values indicate higher specificity of the tool. We randomly subsampled either three or one replicate from the six available replicates, performing nine inter- and intra-condition comparisons, and calculated the IIR by averaging the number of DSEs.

To evaluate each tool’s ability to detect biologically meaningful splicing events in the absence of biological replicates, we analyzed RNA-seq data of one replicate with KD of NMD factors (36) by different tools. We calculated the proportion of up-regulated differentially spliced junctions associated with transcripts labeled as NMD target (Ensembl 106), following the approach used in a previous study (49). Briefly, each annotated intron was classified into one of several categories, based on the following priority: protein_coding > processed_transcript > lncRNA > unprocessed_pseudogene > retained_intron > nonsense_mediated_decay. Only introns uniquely used in transcripts labeled as nonsense_mediated_decay were assigned to that category. In cases where an intron was associated with multiple categories, it was classified according to the highest-priority category in the hierarchy. Note that Whippet is not analyzed here, as the tool does not provide junction-level information associated with the analyzed splicing events, making it unsuitable for this type of analysis.

Additionally, we assessed the correlation between changes in splicing and gene expression. For each gene, we calculated the log 2 base of CPM fold change and the log 2 base of the absolute value of the PSI difference of the alternative splicing event with the strongest change in the gene. We then measured Spearman’s ρ to evaluate the correlation between splicing changes and expression for each tool.

### Data visualization

We used the UCSC Genome Browser (50) to visualize RNA-seq and HITS-CLIP read coverage in regions of interest. All graphs were generated using the pandas (version 1.5.0), matplotlib (version 3.6.1), and seaborn (version 0.12.0) packages in the Python environment (version 3.10.6) in a Singularity container (jupyter/scipy-notebook:notebook-6.4.12 in Docker Hub).

### Data Availability

The raw real RNA-seq data used in this study are available at GEO or SRA with the following IDs: GSE90179, GSE90180, GSE159993, GSE57433, GSE133291, GSE84803, GSE149491, SRP142522, SRP013558, SRP014759, and SRP059357. The sc/snRNA-seq data used in this study is available at GSE116138 and GSE190940. The HITS-CLIP data used in this study is available at GSE103315, GSE39911, and GSE38497.

### Code Availability

Shiba and scShiba codes are available at https://zenodo.org/records/14713893.

## Results

### The Shiba method for alternative splicing analysis

To enable robust and reproducible detection of alternative splicing events using RNA-seq data, we constructed the four-step pipeline illustrated in Figure 1A-B. In the first step, using aligned RNA-seq data and reference gene annotation in GTF format, reference-guided transcript assembly is performed by StringTie2 (23). This step identifies novel transcripts not yet described in the reference gene annotation file, enabling detection of both annotated and unannotated splicing events. Therefore, by utilizing exon body read coverage, Shiba is more sensitive in detecting unannotated events compared to existing methods that rely solely on junction read information.

**Figure 1:**
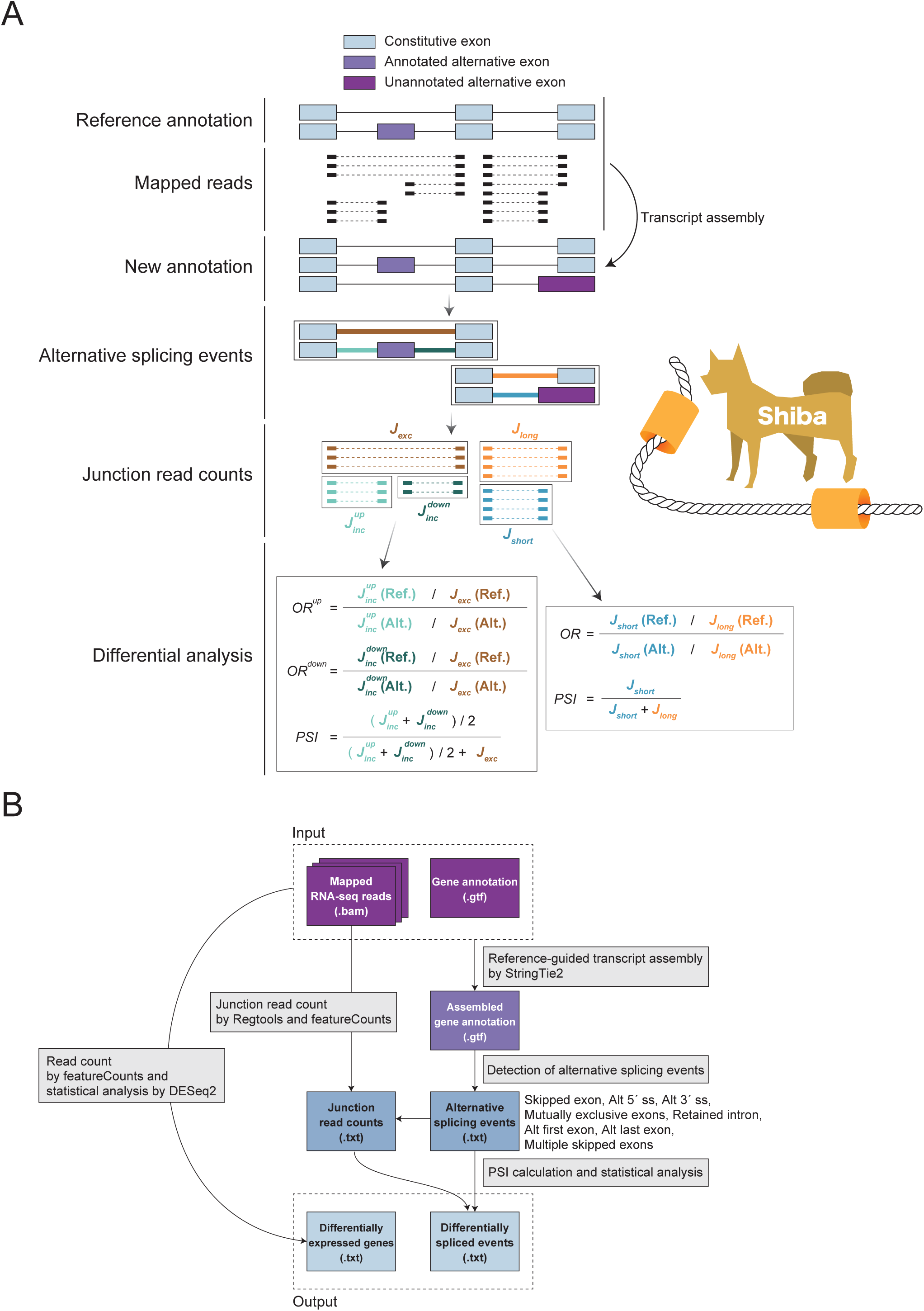
Overview of Shiba. (A) Visualization of Shiba’s statistical analysis process for detecting differentially spliced events. (B) Schematic representation of the Shiba pipeline illustrating the four key steps: reference-guided transcript assembly, identification of alternative splicing events, read counting, and statistical analysis. Shiba output also includes differentially expressed genes.

In the second step, based on the information of transcript structures built in the first step, all alternative splicing events are listed and divided into eight categories: skipped exon (SE), alternative five prime splice site (FIVE), alternative three prime splice site (THREE), mutually exclusive exons (MXE), retained intron (RI), alternative first exon (AFE), alternative last exon (ALE), and multiple skipped exons (MSE). Each event is labeled as “annotated” or “unannotated”. This label can be useful for the functional characterization of cryptic splicing in disease conditions.

In the third step, the counting of reads mapped to each exon-exon and exon-intron junction is performed using RegTools (24) and featureCounts (25). For exon-exon junctions, users can set the anchor length of reads and the maximum and minimum length of the intron to be analyzed. In the final step of the pipeline, the Percent Spliced In (PSI) is calculated and statistical analysis is performed to identify differentially spliced events (DSEs) between the two groups.

To allow the method to be agnostic to sample sizes, including *n*=1 for unique or hard-to-replicate biological samples, individual cell clusters from scRNA-seq, or a single replicate of RNA-seq data (e.g., those published during the early age of RNA-seq), Fisher’s exact test is performed for two-by-two tables of inclusive and exclusive junction read counts between the reference and alternative groups (Supplementary Figure S1). The odds ratio (OR) and *P*-value are then calculated for each alternative splicing event. Junction reads of replicated samples for each group are pooled to gain read coverage.

These values together with the difference in PSI between the two groups (ΔPSI) are used to define DSEs. In the default setting, events meeting all the following criteria will be classified as DSEs: (i) OR is greater than 3/2 or smaller than 2/3, (ii) FDR-adjusted *P* (*q*) is less than 0.05, and (iii) |ΔPSI| is greater than 10. In the case of SE and RI events, two-by-two tables are created separately for the upstream and downstream inclusive exon-exon/exon-intron junctions, resulting in two ORs and two *P*-values (Supplementary Figure S1A). In this scenario, an event is claimed as a DSE only if both ORs exceed the threshold and the larger *q*-value is below 0.05. MXE events have four inclusive exon-exon junctions connecting each of the two flanking constitutive exons with each of the mutually exclusive exons. Four two-by-two tables are created for the calculation of four ORs and *P*-values (Supplementary Figure S1B). In this scenario, an event is detected as a DSE only if all four ORs exceed the threshold and the largest *q*-value is below 0.05. For the FIVE, THREE, AFE, and ALE events, a two-by-two table is created for the two exon-exon junctions corresponding to each alternative splice site to calculate the OR and *P*-value to claim a DSE (Supplementary Figure S1C). In the case of MSE events with *n* skipped events, *n*+1 two-by-two tables are created for each inclusive exon-exon junction, resulting in *n*+1 ORs and *P*-values (Supplementary Figure S1D). An event is claimed as a DSE only if all ORs exceed the threshold and the largest *q*-value is below 0.05.

Pooling reads is an effective strategy to increase read coverage while losing biological variability within a condition. To account for variations between replicates, Shiba implements Welch’s t-test for PSI values of replicates between conditions, a method herein referred to as Shiba+. In this approach, DSEs are identified only if they meet Shiba’s original criteria and the Welch’s t-test *P* is below 0.05. This method enhances the statistical assessment of splicing differences by taking into account the variability present across replicates, leading to more accurate and reliable results.

### Performance evaluation

To assess Shiba, we performed comprehensive comparisons with other highly cited computational tools, including rMATS (4, 5), SUPPA2 (6), Whippet (7), MAJIQ (12, 13), and LeafCutter (14), using both simulated and real RNA-seq datasets. We evaluated the power of Shiba, Shiba+ (Shiba’s original criteria plus Welch’s t-test), rMATS, SUPPA2, MAJIQ het (MAJIQ’s heterogen module quantifying splicing profiles for each sample separately and applying statistical tests to deal with heterogenous samples), MAJIQ deltapsi (MAJIQ’s deltapsi module that assumes a shared splicing profile for a given group of samples and accumulates evidence across samples to infer splicing profiles), and LeafCutter. We applied the computational tools to analyze simulated RNA-seq data. Specifically, we simulated RNA-seq reads to represent alternative splicing events showing differential splicing and those showing unaltered splicing between two groups in the mouse genome (Supplementary Figure S2). We set multiple conditions of read coverages (low, moderate, high, and very high), biological replicates (ranging from 1 to 10), and variations between replicates (low, moderate, and high).

The read coverages of simulation were set to generate junction reads comparable to real RNA-seq data mapped to each splicing event. We confirmed that, in real RNA-seq data, the median number of junction reads mapped to alternative skipped exons was 99 for datasets with 40 million mapped reads (calculated from data registered in GEO under GSE57433) (36) and 329 for those with 100 million mapped reads (calculated from data registered in GEO under GSE149491). In our simulated RNA-seq data, the median number of junction reads mapped to alternative skipped exons was 57 for low, 113 for moderate, 170 for high, and 576 for very high read coverage conditions, closely resembling the range observed in real datasets.

The variation between replicates was set to mimic biological heterogeneity within the same condition. In the low variance condition, individual samples exhibit nearly identical PSI values for each splicing event. For moderate and high variance conditions, we introduced increasing levels of variability, simulating biological differences between replicates within a given condition (Supplementary Figure S3A-B). Since SUPPA2 does not support differential splicing detection using one replicate data, it was excluded from the 1 vs. 1 comparison.

Each combination of read coverage, replicate number, and variation between replicates was analyzed to evaluate its power of differential splicing detection. In addition to annotated splicing events, we included unannotated splicing events in the simulation by removing records of inclusion isoforms from the reference gene annotation file. We evaluated the performance across methods of detecting differential splicing for five classical splicing events (SE, FIVE, THREE, MXE, and RI), using metrics of false negative rate (FNR), false positive rate (FPR), and Matthews correlation coefficient (MCC).

Shiba consistently demonstrates the lowest FNR across almost all conditions, indicating superior sensitivity in detecting DSEs, both for annotated and unannotated splicing events (Supplementary Figure S4). Shiba’s lower FNR compared to others stands out the most in cases of moderate or high variance, and particularly in combination with low read coverage. Shiba’s advantage is more apparent in scenarios of fewer replicates. Shiba+, MAJIQ het, and LeafCutter are in the second best group with slightly higher FNR than Shiba. This group displays decreasing FNRs with increasing replicate numbers, narrowing their difference with Shiba. Other methods, such as rMATS, SUPPA2, and Whippet, had higher FNRs, particularly under high variance conditions, though rMATS showed notable improvement with increasing replicates. MAJIQ deltapsi has the highest FNRs for annotated splicing events. SUPPA2 and Whippet demonstrated the highest FNRs for unannotated splicing events, which is expected as these tools are not designed to detect unannotated splicing events. The differences between these tools persist in low to high read coverage.

At very high read coverage, all tools, except for MAJIQ deltapsi, continued to reduce FNR. Shiba consistently demonstrated the lowest FNR for annotated event detection, outperforming other methods. Shiba+, MAJIQ het, and LeafCutter performed similarly well. rMATS showed improvement with read coverage but exhibited a still higher FNR than Shiba. For unannotated event detection, Shiba ranked the second and MAJIQ het achieved the lowest FNR, particularly when the number of replicates increased. While SUPPA2 and Whippet also improved under higher read coverage, they continued to exhibit very high FNR in unannotated event detection.

In summary, Shiba consistently showed the lowest FNR. In conditions of low replicates, high variance, and lower read coverage, it demonstrated exceptional sensitivity in detecting DSEs. Shiba+, while not as sensitive as Shiba in terms of FNR, performed similarly to MAJIQ het and LeafCutter across all conditions.

FPR is generally comparable between all tools (Supplementary Figure S5). Whippet showed the highest FPR in conditions of low replicate number and low variance. Shiba exhibited a slightly higher FPR under conditions of high variance or a combination of low replicate numbers and moderate variance. Nevertheless, Shiba+ substantially reduced Shiba’s FPR to levels indistinguishable to those of rMATS, SUPPA2, MAJIQ het, MAJIQ deltapsi, and LeafCutter. As the number of replicates and read coverage increased, all tools perform satisfactorily well. Note that this simulation did not mimic junction read imbalance in real RNA-seq data, resulting in low FPRs derived from all tools. Therefore, FPR from this simulation does not necessarily provide sufficient evidence to distinguish these methods.

Next, we assessed the performance of each tool using MCC, a balanced metric that considers true positives, true negatives, false positives, and false negatives (Figure 2A-B). Shiba consistently performed well, maintaining higher MCC values across all conditions. The second best group includes Shiba+, MAJIQ het, and LeafCutter that displayed similarly strong MCC values. For detecting unannotated splicing events MAJIQ het edge Shiba under the conditions of very high read coverage or low variance, but Shiba still performed better under more scenarios. rMATS showed moderate performance, improving only with more replicates and higher read coverage, and consistently lagged the top-performing tools. SUPPA2, Whippet, and MAJIQ deltapsi showed lower MCC values than others across all conditions, particularly in high variance settings and with fewer replicates, indicating less balanced performance in terms of false positives and false negatives. These findings highlight Shiba as a top performer for detecting DSEs, particularly in scenarios with low and moderate read coverage and fewer replicates. On the other hand, Shiba+, LeafCutter, and MAJIQ het were particularly effective under conditions of greater biological variance, demonstrating their strength in handling complex data variability.

**Figure 2:**
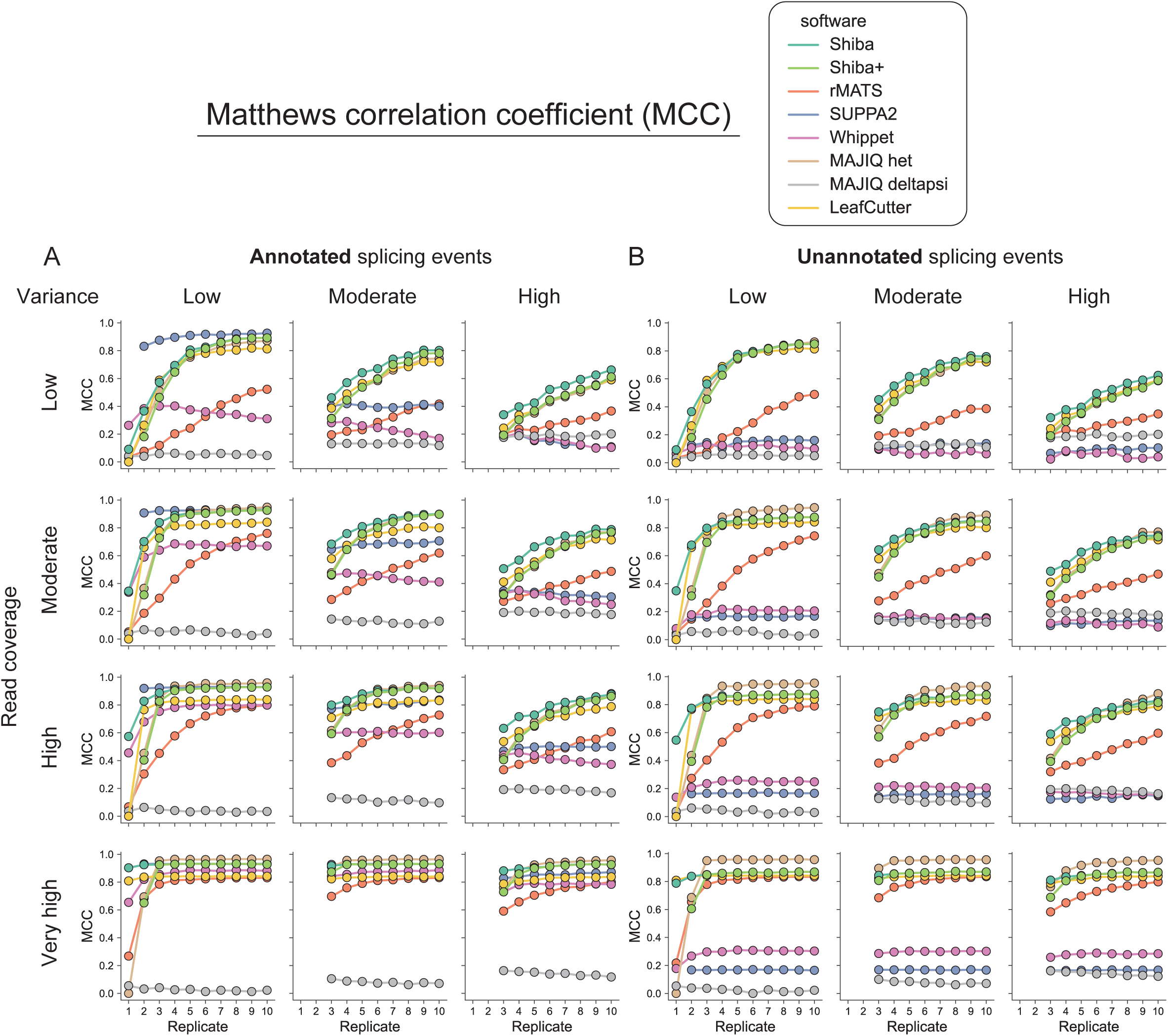
Performance evaluation on detecting differentially spliced genes in simulated RNA-seq data. (A) Matthews correlation coefficient (MCC) for annotated splicing events detection by eight computational tools/modules (Shiba, Shiba+, rMATS, SUPPA2, Whippet, MAJIQ het, MAJIQ deltapsi, and LeafCutter) in different combinations of sample variations (low, moderate, and high), read coverages (low, moderate, high, and very high), and biological replicates (ranging from 1 to 10). (B) The corresponding metric for unannotated splicing events detection.

In addition to evaluating performance for these five classical splicing events, we conducted the same analysis for alternative first exons and last exons (AFE and ALE) (Supplementary Figure S6, S7, S8), and multiple skipped exons (MSE) (Supplementary Figure S9, S10, S11). The performance results were consistent with those observed for the classical events. Shiba maintained its strong performance across all conditions, demonstrating high MCC values and remaining a top performer. These findings demonstrate that Shiba remains highly effective for detecting a diverse range of splicing events, including AFE, ALE, and MSE, in both annotated and unannotated scenarios. Moreover, the consistent performance of Shiba and Shiba+ across various event types and conditions underscores their utility in analyzing complex RNA-seq datasets, particularly in situations where read coverage is limited or biological variance is high.

### Evaluation of robustness from sampling biological replicates

A robust computational tool, when applied to detect DSEs, should show greater consistency when sampling subsets of biological replicates. To compare different tools, we assessed their performance using real dataset with many replicates: RNA-seq samples from mouse adult liver and kidney tissues (32). The six biological replicates allowed us to scramble the replicate combinations to assess reproducibility, employing the reproducibility ratio (RR) metric (13). In essence, RR quantifies the proportion of DSEs identified by an algorithm in dataset D that is consistently identified using a similar replicate dataset D’. Our RR analysis subsampled three or one out of six biological replicates from each condition (liver or kidney), resulting in nine instances of differential splicing analysis.

In both 3 vs. 3 (Figure 3A) and 1 vs. 1 scenarios (Figure 3B), Shiba demonstrates stably high RR values across a wide range of top *n* events, indicating its robustness and consistency in detecting DSEs. The only other method showing stable performance in the 3 vs. 3 scenario is LeafCutter, but with substantial lower RR values than those of Shiba. MAJIQ het displays the least RR values among all. All other methods can exhibit similar or even greater RR values in certain windows of event calling, but clearly lack Shiba’s consistency. For example, rMATS, SUPPA2, and Whippet perform substantially worse when smaller numbers of events are called. The superior performance of Shiba is even more obvious in the 1 vs. 1 scenario. Shiba again displays a stably high RR, maintaining a high fraction of reproducible events even under minimal replicate conditions. This demonstrates Shiba’s strength in scenarios with limited sample size, ensuring high reproducibility and reliable performance when it is challenging to obtain biological replicates.

**Figure 3:**
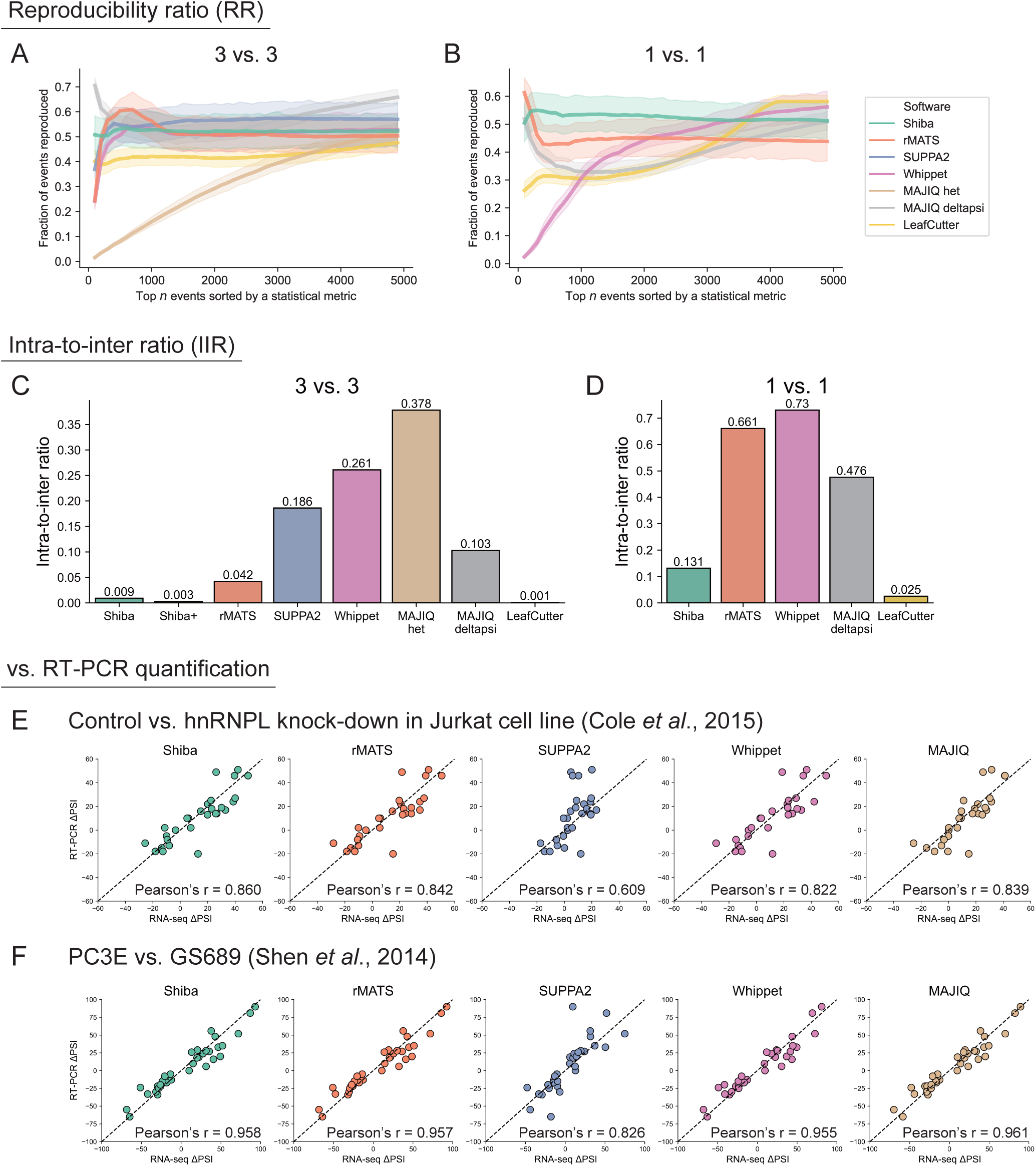
Performance evaluation using real RNA-seq data. (A, B) The reproducibility ratio (RR) metric when subsampling three (A) or one (B) out of six biological replicates in real RNA-seq samples from mouse adult liver and kidney (32). Events are ranked by a statistical metric with a 95% confidence interval (see Methods). The X-axis represents the ranked number of events reported, while the Y-axis indicates the fraction of those events reproduced among the same number of top-ranking events in repeated analyses using different sample replicates. (C, D) The intra-to-inter ratio (IIR) metric for group size of three (C) and one (D). The IIR is ratio between the number of differentially spliced events derived from comparing the same tissue (liver vs. liver) and the number of events derived from comparing the different tissues (liver vs. kidney). (E, F) Correlation of ΔPSI values in RNA-seq and RT-PCR of samples from (E) control vs. hnRNPL knockdown in Jurkat cell lines (35) and (F) PC3E vs. GS689 cell lines (4).

To further evaluate the specificity of the tool performance, we calculated the intra-to-inter ratio (IIR) (13). IIR is an approximation of false discovery proportion. The IIR is obtained by comparing the number of DSEs identified in intra-condition comparisons (e.g., liver vs. liver) to those identified in inter-condition comparisons (e.g., liver vs. kidney). Since DSEs detected in intra-condition comparisons are more likely to be false positives, a lower IIR value indicates lower type I errors or greater specificity of the tool.

In the 3 vs. 3 scenario (Figure 3C), LeafCutter, Shiba, and Shiba+ shows the lowest IIR values (0.001, 0.009, and 0.003, respectively), an order lower than results from other tools. This supports the notion that Shiba and Shiba+ not only maintain high reproducibility but also accurately identify true DSEs with minimal false positives. MAJIQ het shows the highest IIR value (0.378), suggesting a higher occurrence of false positives and therefore lower specificity compared to the other tools.

In the 1 vs. 1 scenario (Figure 3D), LeafCutter again achieves the lowest IIR value (0.025). Shiba ranks second with a low IIR of 0.131, reinforcing its ability to maintain specificity under conditions with few biological replicates. This consistency highlights Shiba’s adaptability and reliability across different replicate configurations. In contrast, Whippet, rMATS, and MAJIQ deltapsi show much higher IIR values (0.73, 0.661, and 0.476, respectively) in the 1 vs. 1 setting, showing that they are not suitable for analysis with very limited sample numbers.

In summary, the IIR analysis confirms that Shiba maintains both high reproducibility and specificity across different replicate configurations, making it a reliable tool for DSE detection even in challenging scenarios with limited sample size. Importantly, Shiba effectively balances both reproducibility and specificity. Whippet, rMATS, and MAJIQ deltapsi, although competitive in certain conditions, show reduced specificity, particularly when fewer replicates are available.

### Evaluation of accuracy in ΔPSI quantification

To assess the accuracy of each tool in quantifying exon-level splicing changes, we compared the RNA-seq-derived ΔPSI values with those obtained from RT-PCR experiments (gold standard). We found two published datasets that contain both RNA-seq and PSI values of RT-PCR results: (i) samples of two prostate cancer cell lines, PC3E and GS689, which were used by the rMATS developers to validate rMATS results (4), and (ii) cultured Jurkat cell line with *hnRNPL* knockdown (KD) (35), used by Vaquero-Garcia et al. to validate MAJIQ results (12). Pearson’s correlation coefficient (r) was calculated for each tool to quantify the strength and direction of the relationship between RNA-seq-based and RT-PCR-derived ΔPSI values. Note that LeafCutter lacks the ability to estimate exon-level PSIs, making it unsuitable for this type of analysis.

Shiba ranked the best in the *hnRNPL* KD dataset (Figure. 3E). Note that the RT-PCR results in this study represent a selective set of exons to confirm the MAJIQ analyses, but Shiba still outperform MAJIQ (Pearson’s r = 0.86 vs 0.839). The RT-PCR results in the PC3E vs. GS689 study were used to support the rMATS analysis, and Shiba exhibited a slightly higher Pearson’s coefficient than rMATS (Figure. 3F). MAJIQ performed the best with this dataset. Overall, Shiba outperformed rMATS, SUPPA2, and Whippet in both datasets. SUPPA2 displayed a noticeably lower correlation than others, suggesting some discrepancies in its ΔPSI estimates.

### Shiba considers junction read imbalance to reduce false positives

Shiba employs Fisher’s exact test on both sides of inclusive junctions. This careful approach allows Shiba to accurately assess and account for previously uncharacterized junction read imbalance, minimizing false positives. Our analysis of the Yeom et al. dataset (34) revealed that 30.71% of alternative skipped exons exhibit a >2-fold difference in read counts between upstream and downstream inclusion junctions (Figure 4A-B), demonstrating the prevalence of junction read imbalance in real RNA-seq data. We hypothesized that the higher IIR values and a larger number of false positives derived from existing computational tools (Figure 3C-D) are partly due to inadequate consideration of junction read imbalance.

**Figure 4:**
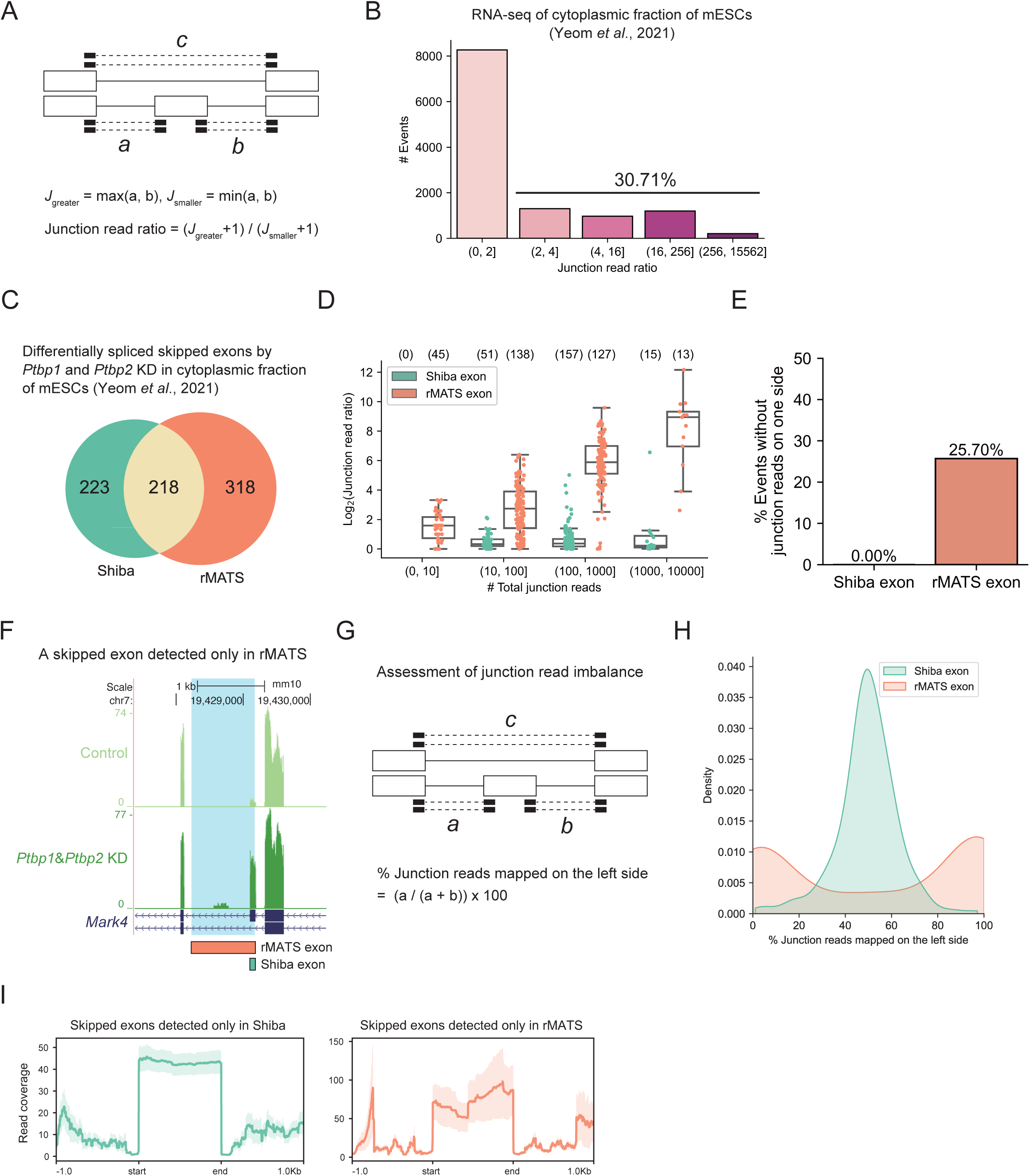
Shiba considers junction read imbalance. (A) Schematic representation of junction read ratio in a skipped exon (SE). (B) Number of SE events for each bin of junction read ratio (0–2, 2–4, 4–16, 16–256, and 256–15,562) in RNA-seq data of cytoplasmic fraction of mESCs (34). (C) Comparison of DSE events detected by Shiba vs. rMATS using the cytoplasmic fraction of mESC with siRNA-mediated double knockdown of Ptbp1 and Ptbp2 (34). There are 218 events common to both tools, 223 additional events unique to Shiba, and 318 events exclusive to rMATS. (D) Junction read ratios by total junction read counts in four bins (0–10, 10–100, 100–1000, and 1000–10,000) for events exclusively identified by Shiba and rMATS. Numbers above each category represent the sample size. (E) Percentage of events with no reads on one side of junctions for events exclusively identified by Shiba and rMATS. (F) A specific SE event in Mark4 gene, exclusively identified by rMATS, shows a potential error in exon definition as illustrated by the discrepancy from the RNA-seq browser tracks. The exons exclusively identified by rMATS and Shiba are shown in the orange and green colored boxes, respectively, below the gene annotation. (G) Schematic showing a measurement of junction read imbalance. (H) Distribution of the percentage of junction reads mapped on the left side for events exclusively identified by Shiba and rMATS. (I) Aggregation plots of read coverage on the meta-exon, demonstrating even distribution for Shiba-identified events and uneven distribution for rMATS-identified events.

To test this hypothesis, we compared the DSEs detected by Shiba vs. the popular rMATS using the Yeom et al. dataset, i.e., the cytoplasmic fraction of mESCs upon siRNA-mediated double KD of *Ptbp1* and *Ptbp2* (34). This comparison revealed 218 differentially spliced skipped exons detected by both tools, with Shiba identifying an additional 223 events and rMATS detecting 318 events exclusively (Figure 4C). We then determined the junction read ratios for exon groups categorized by their number of total junction reads. Our analysis revealed that regardless of total junction read numbers, rMATS-identified events exhibited substantially higher junction read ratios than events exclusively identified by Shiba (Figure 4D).

Moreover, we observed that junction read imbalance tended to increase with the total junction read count for rMATS-identified events. Therefore, the high junction read ratios for rMATS events were not due to low coverage. In contrast, the junction read ratios remained steadily low for Shiba-identified events, showing Shiba’s robustness against such biases (Figure 4D). Interestingly, about the same numbers of Shiba exons and rMATS exons contain (1000, 10000] junction reads yet exhibiting completely different junction read ratios. Similar observation was made for the exon group with (100, 1000] junction reads, indicating that the two algorithms can vary in exon definition.

Remarkably, 25.70% of the rMATS-identified events contained no reads at all on one side of the junctions, contrasting with events exclusively identified by Shiba which all had at least one read on both sides of the junctions (Figure 4E). The absence of junction reads in some of these rMATS-identified events is associated with erroneous junction site determination, illustrated by a DSE claimed by rMATS in *Mark4* gene (Figure 4F). In this case, the rMATS exon and Shiba exon share the common upstream 3’ splice site but differ in their downstream 5’ splice site, where no junction reads support the 5’ splice site of the rMATS exon. Genome browser tracks of the RNA-seq data clearly agree with the Shiba results. These kinds of “imbalanced” alternative skip exons were not uncommon in the rMATS output (Supplementary Figure S12).

Next, we quantitatively evaluated junction read imbalance by computing the percentage of junction reads mapped on either side (Figure 4G). Our analysis unveiled a bimodal distribution in the metric for events exclusively identified by rMATS, indicating that, in the majority of these events, the number of junction reads on one side greatly differed from those on the other side. In contrast, the metric for events exclusively identified by Shiba exhibited a normal distribution with an average score of nearly 50%, indicating a balanced distribution of junction reads on both sides of the exons for these events (Figure 4H). This result suggests that rMATS tends to identify events with a notable junction read imbalance. The aggregation plots of read coverage also demonstrate that events exclusively identified by Shiba show an even distribution of read coverage on the meta exonic region, while rMATS-identified events exhibit uneven read distribution despite overall higher read coverage (Figure 4I). The identification of DSEs with imbalanced junction reads by rMATS is not limited to cassette exons but also observed in all other types of alternative splicing events (Supplementary Figure S13).

Taken together, these data underscore the importance of considering read coverage patterns and uniformity in addition to coverage depth for accurate event identification and quantification (51). Shiba’s ability to separately consider the two junctions increases reliability of the splicing quantification and minimizes erroneous calling of DSEs.

### Evaluation of runtime

We evaluated the runtime of each tool using real RNA-seq data from the Yeom et al (34) (Supplementary Figure S14). We allocated 16 CPUs for Shiba, rMATS, MAJIQ, LeafCutter, STAR, and Salmon. SUPPA2 and Whippet do not support multi-core usage, so they were executed on a single CPU. Additionally, we evaluated the runtime of SnakeShiba, which is a Snakemake-based and faster version of the Shiba pipeline. We found Shiba, SnakeShiba, rMATS, MAJIQ, and LeafCutter, utilizing the mapped files as input, took 56.6, 45.9, 37.9, 42.1, and 42.1 minutes, respectively. Shiba implements DESeq2 to perform a differential gene expression analysis that the other tools do not do (Figure 1A), resulting in a slightly longer runtime. SUPPA2, including the expression quantification step using Salmon, required 20.1 minutes. Whippet, which internally performs pseudo-mapping, took 75.4 minutes. Overall, the time difference is negligible against the backdrop of time needed for data acquisition. In summary, Shiba is capable of evaluating transcriptome data for both DEG and DSE in a multifaceted and speedy manner.

### Shiba analysis of single-replicate RNA-seq data

A notable strength of Shiba is its applicability to RNA-seq data without biological/technical replicates. In the early days of RNA-seq, biological samples often lacked replicates. Even nowadays, while three biological replicates represent the experimental gold standard, they are inherently insufficient for regression statistics. We compared Shiba with other tools that can perform differential splicing analysis of single-replicate data (rMATS, Whippet, MAJIQ deltapsi, and LeafCutter). To unbiasedly assess their performance, we seek a single-replicate dataset in which inhibition of nonsense-mediated mRNA decay (NMD) leads to changes in mRNA abundance and correlated changes in steady-state exon inclusion ratios for alternative splicing events coupled with NMD (AS-NMD). As such, the performance of DSE calling can be evaluated by the degree of correlation between ΔPSI and gene expression changes.

We found RNA-seq data of double KD of *XRN1* and *UPF1* for its absence of replicates (36) (Figure 5A). XRN1, a cytoplasmic 5′–3′ exonuclease, plays a pivotal role in the swift removal of decay intermediates (52), and UPF1 serves as a key component in nonsense-mediated decay (NMD) (53). Lykke-Andersen et al. showed in their study that double KD of *XRN1* and *UPF1* effectively inhibits NMD (36). Each tool identified distinct sets of differentially spliced genes (DSGs), yielding a total of 9,707 DSGs (Figure 5B). Shiba identified 1,516 DSGs in total, more than MAJIQ and LeafCutter but substantially less than rMATS and Whippet. Interestingly, only 259 DSGs, or 2.7% of total DSGs, were identified by all tools, illustrating significant variability among different methods.

**Figure 5:**
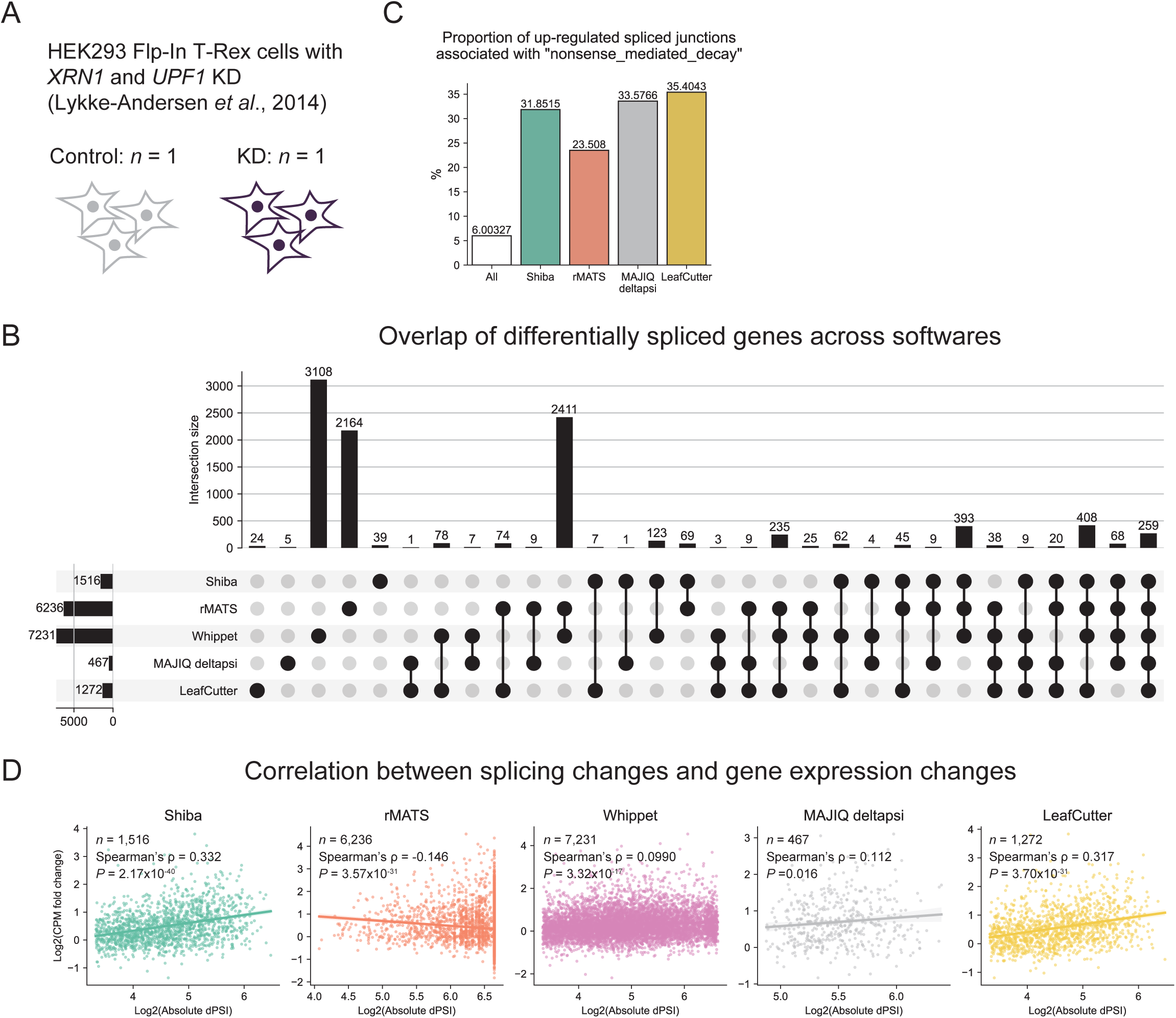
Shiba analysis of single replicate RNA-seq data. (A) Experimental design of RNA-seq for HEK293 Flp-In T-Rex cells with siRNA-mediated double knockdown of XRN1 and UPF1, lacking replicates (36). (B) An upset plot of overlap of differentially spliced genes across Shiba, rMATS, Whippet, MAJIQ deltapsi, and LeafCutter. (C) Proportion of up-regulated spliced junctions associated with transcripts labeled as “nonsense_mediated_decay”. (D) Correlation between splicing changes and gene expression changes for events identified by each software. The X-axis represents the log 2 base of the absolute value of the PSI difference for the detected DSEs. The Y-axis represents the log 2 base of CPM fold change of the corresponding gene. If a gene has multiple DSEs, the splicing event exhibiting the largest difference is selected.

To assess whether the DSG calling globally agrees with annotation of AS-NMD events, we calculated the proportion of up-regulated differentially spliced junctions associated with annotated NMD transcripts. NMD-sensitive alternative transcripts are expected to be up-regulated upon NMD inhibition, leading to an increase in the splice junction specific to the NMD-sensitive transcripts, i.e., the inclusive junctions associated with inclusion-induced NMD transcripts, and the skipping junctions associated with exclusion-induced NMD transcripts. Whippet results lack the junction information associated with the analyzed splicing events so are unsuitable for this type of analysis. Our analysis showed that that 31.8% of the up-regulated junctions identified by Shiba are NMD-associated junctions, substantially higher than the proportion of AS-NMD transcripts among all transcripts (6.00%) (Figure 5C). MAJIQ deltapsi (33.58%) and LeafCutter (35.40%) yielded similar results, indicating that the three are similarly proficient in identifying AS-NMD events. In contrast, up-regulated junctions identified by rMATS have obviously less NMD-associated junctions (23.51%), indicating that as an exon-centric splicing calling method rMATS is not as adept as Shiba in handling single-replicate RNA-seq data.

To further examine the performance of DSE calling upon NMD inhibition, we calculated the correlation between changes in steady-state inclusion ratios (absolute PSI difference) and gene expression changes (log2 CPM fold change) for each tool. This correlation tests whether the identified DSEs are coupled with changes in steady-state mRNA abundance, as would be expected for AS-NMD targets. As shown in Figure 5D, Shiba exhibited a positive correlation (Spearman’s ρ = 0.332, *P* = 2.17×10^-40^), confirming that the splicing events detected by Shiba are indeed globally associated with changes in gene expression. LeafCutter demonstrated positive correlation with a slightly weaker correlation coefficient and p-value (Spearman’s ρ = 0.317, *P* = 3.70×10^-31^) than those of Shiba. By contrast, rMATS, MAJIQ deltapsi, and Whippet showed much lower or even negative correlations (ρ = −0.146, 0.112, and 0.099, respectively).

Overall, these analyses support the strength of Shiba in accurately identifying DSEs in the absence of replicates. In the NMD inhibition dataset, Shiba detects a broad range of DSGs whose steady-state abundance are driven by changes in NMD sensitive isoforms, thereby reflected by changes in their splicing ratios. Therefore, Shiba’s capacity to extract biologically relevant splicing regulation from RNA-seq data of limited replications underscores its broad utility in mining various data resources for novel biological insights.

### Application of scShiba on single-cell/nucleus RNA-seq data to define differential splicing between cell groups

Shiba application on one-replicate data presents an opportunity to derive differential splicing from sc/snRNA-seq data with confident statistical metrics. Toward this goal, we developed scShiba, an alternative pipeline of Shiba for sc/snRNA-seq data that extends the capabilities of Shiba to sc/snRNA-seq data, allowing for a comprehensive analysis of alternative splicing dynamics between cell clusters. Due to the limited read coverage for individual cells, a fundamental strategy employed by the Shiba method involves aggregating reads for samples of interest, e.g., the same cell type or a cell cluster. This pooling approach for DSE detection enhances our ability to derive novel insights from sc/snRNA-seq data. scShiba takes several output files of STARsolo (28), a text file of user-defined cell type annotations, and a gene annotation file (Figure 6A). The other scShiba steps are identical to those of Shiba: pooling junction reads for each cell type, detecting alternative splicing events, calculating PSI, and performing statistical tests.

**Figure 6:**
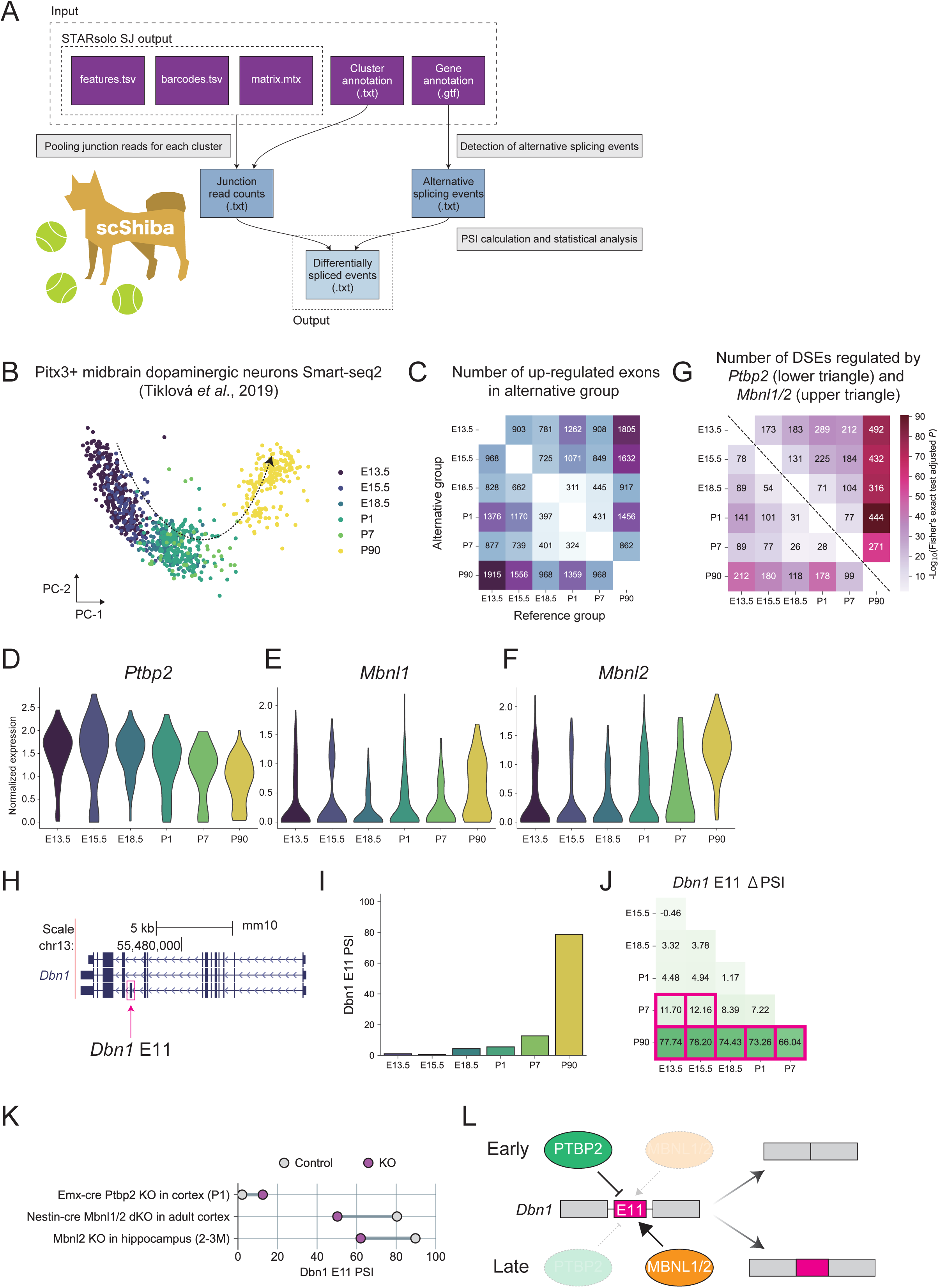
scShiba identifies neuronal-stage-specific alternative splicing events using single-cell RNA-seq data. (A) Workflow of scShiba, an extension of Shiba designed for single-cell/nucleus (sc/sn) RNA-seq data. (B) Smart-seq2 scRNA-seq data from mouse Pitx3+ midbrain dopaminergic neurons (41), plotted by the first two principal components of the normalized gene expression matrix. (C) A heatmap of numbers of differentially spliced events (DSEs) detected in scShiba. The number in each box represents up-regulated DSEs (q < 0.05, Δ PSI > 10) in the alternative group compared to the reference group. (D-F) Expression levels of Ptbp2 (D), Mbnl1 (E), and Mbnl2 (F) for each cell type in the dopaminergic neurons at different ages. (G) A heatmap of the numbers of developmental DSEs that are differentially spliced in Ptbp2 knockout (lower triangle) and Mbnl1/2 knockout (upper triangle). The color represents adjusted P-values of Fisher’ s exact test. (H) Genome browser track of Dbn1 gene and its alternative exon 11 (E11). (I) PSI values of Dbn1 E11 at different stages from the Smart-seq2 data. (J) ΔPSI values of Dbn1 E11 in each comparison between stages. Magenta boxes represent statistical significance (q < 0.05, IΔPSII > 10). (K) PSI values of Dbn1 E11 in control and knockout (KO) samples of Emx1-cre Ptbp2 KO mouse cortices (postnatal day 1) (38), Nestin-cre Mbnl1 and Mbnl2 double KO mouse adult cortices (39), and Mbnl2 KO mouse adult hippocampi (2-3 months old) (40). (L) A model of alternative splicing program of Dbn1 E11 in early and late stages of neuronal maturation.

To test scShiba performance, we analyzed scRNA-seq datasets of differentiating dopaminergic neurons, because developmental splicing regulation of this neuronal subtype had not been examined before. Specifically, we analyzed Smart-seq2 data of mouse Pitx3+ midbrain dopaminergic neurons at embryonic day 13.5 (E13.5), 15.5 (E15.5), 18.5 (E18.5), postnatal day 1 (P1), 7 (P7), and 90 (P90) (41), aiming to identify DSEs during neuronal maturation (Figure 6B). Our scShiba analysis revealed significant differential splicing across each time point, identifying a total of 28,866 DSEs (*q* < 0.05, |ΔPSI| > 10) (Figure 6C). Next, we explored possible upstream RBP regulators. RNA binding protein PTBP2 regulates alternative splicing during early differentiation of cortical neurons and motor neurons (54–56). In later stages of cortical brain development, other RBPs like MBNL1 and MBNL2 play significant roles in alternative splicing regulation of synaptic proteins (39). These observations were previously derived from bulk RNA-seq data. Using these DSEs, we examined whether dopaminergic neuron differentiation is similarly mediated by PTBP2 and MBNL1/2. We found high *Ptbp2* expression in early differentiation stages of dopaminergic neurons (Figure 6D) and increased expression of *Mbnl1* and *Mbnl2* in later stages (Figure 6E-F). We first determined splicing events regulated by these RBPs using RNA-seq samples from Emx1-cre *Ptbp2* knockout (KO) mouse cortices (postnatal day 1) (38), Nestin-cre *Mbnl1* and *Mbnl2* double KO mouse adult cortices (39), and *Mbnl2* KO mouse adult hippocampi (2–3 months old) (40). We found PTBP2 and MBNL1/2 targets significantly overlapped with DSEs during differentiation and maturation of dopaminergic neurons in all comparisons between developmental stages (Figure 6G).

We categorize regulated exons based on their dual responses to the *Ptbp2* and *Mbnl* KOs and divided them into four groups (Supplementary Figure S15A). The largest group are those up-regulated in *Ptbp2* KO and down-regulated in *Mbnl1*/*2* dKO (Group 2 containing 67 DSEs), suggesting combinatorial regulation with which early PTBP2 expression represses neuronal splicing and late MBNL1/2 expression promotes neuronal splicing. We found 34 of 67 events were DSEs of differentiating dopaminergic neurons from our scShiba analysis. One of these events is a skipped exon in the *Dbn1* gene (*Dbn1* E11), encoding drebrin, a post-synaptic actin-binding protein (57) (Figure 6H). scShiba shows the PSI values of this exon were below 15 during early neuronal differentiation stages (E13.5 ∼ P7), and rose to around 80 at the later stage (P90) (Figure 6I), resulting in a statistically significant difference of PSIs in the comparisons of E13.5 vs. P7, E15.5 vs. P7, and P90 vs. others (Figure 6J). This is consistent with previous study showing that the E11+ isoform of *Dbn1* is predominant in the adult whole brain (57), demonstrating a developmental regulation universal to different neuronal subtypes. E11 is up-regulated in *Ptbp2* KO (PSI in KO = 12.54, PSI in control = 2.10, ΔPSI = 10.44, *q* = 1.71×10^-151^) and down-regulated in *Mbnl1* and *Mbnl2* double KO (PSI in KO = 50.27, PSI in control = 80.38, ΔPSI = 30.11, *q* = 1.97×10^-61^) and *Mbnl2* KO (PSI in KO = 62.05, PSI in control = 89.70, ΔPSI = 27.65, *q* = 2.95×10^-33^) (Figure 6K). We further analyzed HITS-CLIP data of mouse brain tissues and identified prominent peaks of PTBP2 (46), MBNL1 (47), and MBNL2 (40) in *Dbn1* E11 or downstream introns (Supplementary Figure S15B), providing evidence of direct regulation of this splicing event by PTBP2 and MBNL1/2. These findings indicate an alternative splicing regulatory mechanism of synaptic protein *Dbn1* during neuronal maturation shared between cortical and dopaminergic neurons: PTBP2 repression during early development and MBNL1/2 promotion of inclusion during later stages (Figure 6L).

To further test scShiba’s utility in differential splicing analysis, we analyzed 10x snRNA-seq data from the mouse primary visual cortex on postnatal day 28 (43) to investigate DSEs between distinct cell types (Supplementary Figure S16A). Despite the low sequencing depth for each cell and highly biased 3’end sequencing of 10x dataset, scShiba identified a total of 224 DSEs (*q* < 0.05, |ΔPSI| > 10) (Supplementary Figure S16B). For example, we identified a skipped exon in the *Ppp1r12a* gene (*Ppp1r12a* E14) that exhibited differential splicing between excitatory and inhibitory neurons (Supplementary Figure S16C). The splicing levels of this exon were significantly different in excitatory neurons (PSI = 35.8) vs. in inhibitory neurons (PSI = 90.9), resulting in a ΔPSI of 55.2 (*q* = 7.03×10^-16^) (Supplementary Figure S16D-E). This observation aligns with Shiba analyses of bulk RNA-seq dataset of neuronal subtypes (37), in which the PSIs of this exon in various types of excitatory neurons are about 40, significantly lower than those (about 80) in inhibitory neurons (Supplementary Figure S16F).

To infer the underlying regulation of *Ppp1r12a* E14, we examined MBNL expression, because it was suggested to regulate differential splicing between glutamatergic and GABAergic neurons (58). Analysis of 10x snRNA-seq data from the visual cortex unveiled higher expression levels of *Mbnl2* in excitatory neurons compared to inhibitory neurons, while *Mbnl1* showed no disparity between neuronal subtypes (Supplementary Figure S16G-H), indicating that MBNL2 but not MBNL1 is the distinguishing factor. Our Shiba analyses of RNA-seq samples from *Mbnl1/2* KO brains showed MBNL repression of *Ppp1r12a* E14. Its inclusion is up-regulated in *Mbnl1* and *Mbnl2* double KO (PSI in KO = 99.57, PSI in control = 61.24, ΔPSI = 38.33, *q* = 3.31×10^-54^) and *Mbnl2* single KO (PSI in KO = 92.75, PSI in control = 53.15, ΔPSI = 39.60, *q* = 1.11×10^-4^) (Supplementary Figure S16I). Additionally, HITS-CLIP analysis revealed binding signals for both MBNL1 and MBNL2 around *Ppp1r12a* E14, providing evidence of direct regulation (Supplementary Figure S16J). These findings suggest a potential regulatory mechanism for alternative splicing of *Ppp1r12a* E14 in two distinct neuronal subtypes and implicate higher expression of MBNL2 exerting stronger repression of its inclusion in excitatory neurons (Supplementary Figure S16K).

These results, taken together, underscore the utility of scShiba in detecting DSEs using scRNA-seq data from both the Smart-seq and droplet-based (e.g. 10x) technologies. Shiba and scShiba extend the values of scRNA-seq data for systematic splicing analyses and allow cross-platform corroboration with bulk RNA-seq data to derive high-confidence splicing regulation, presenting a valuable tool for unraveling the complexity of alternative splicing in diverse biological systems.

## Discussion

In this study, we developed and conducted an extensive evaluation of the Shiba method for detecting differential alternative splicing from bulk and single-cell/nucleus RNA-seq datasets. The comprehensive comparison with commonly used computational tools, including rMATS, SUPPA2, Whippet, MAJIQ, and LeafCutter, reveals Shiba’s superior well-rounded performance in multiple metrics across various scenarios and read coverages (Supplementary Table). Besides its high sensitivity, Shiba’s strength lies in its accurate detection, even in analyzing data of very limited biological replicates, of both annotated and unannotated DSEs; the latter are commonly overlooked by other popular software such as SUPPA2.

Shiba’s reliability, thanks to meticulous consideration of junction read imbalance for SE, MXE, and RI events, reduces false positives and provides high-confidence DSEs in real RNA-seq data analysis. We observed that the specificity evaluation results differed between simulated and real RNA-seq datasets across tested software. In simulated data, all tools have consistently low FPRs that are almost indistinguishable (Supplementary Figure S5). For real RNA-seq data, tools such as rMATS, Whippet, and MAJIQ showed a notably high IIR (Figure 3C-D), a proximation of false discovery proportion in real data. The discrepancy should be due to junction read imbalance and other FPR-inducing artifacts that are not present in simulated datasets. We therefore weigh the real data evaluation more informatively than simulation. These findings highlight the importance of benchmarking splicing analysis tools with real RNA-seq data to ensure their practical reliability and robustness in addressing experimental noise and read imbalance.

Shiba employs a statistics framework agnostic to sample sizes and can effectively handle *n*=1 datasets (Figure 5). This alleviates a common challenge for experimenters to produce a larger number of biological replicates in adopting the existing computational pipelines. This does not mean we advocate a low number of biological replicates. It only showcases the practical value of Shiba in common experimental settings where many biological replicates are infeasible. Increasing the sample size only increases the statistical power employed by Shiba.

The application of this statistics framework by scShiba resolves a significant challenge in deriving splicing information from scRNA-seq data. Despite the significant impact of single cell sequencing technologies on dissecting the intricacies of gene expression dynamics at a cellular resolution, the increasing volume of scRNA-seq data remain underexplored for splicing dynamics due to low and non-uniform read coverage. Pooling reads from a cell cluster presents a *n*=1 scenario, and scShiba stands out for its verified statistical approaches. scShiba’s ability to uncover splicing variations between cell subtypes or states opens new explorations of the complex context-dependent splicing landscape, including rare splicing events that may be obscured in bulk analyses.

scShiba’s application is not limited to cell clusters or pooling reads of scRNA-seq. When sequencing technology advances to a point where sufficient read coverage can be consistently obtained at the single-cell level, the strength of scShiba to perform differential splicing analysis on single-replicate samples is even more critical, because every single cell is essentially *n*=1 without replicates. scShiba’s capability will allow researchers to fully exploit high-resolution single-cell data, accurately capturing splicing dynamics in individual cells and the full heterogeneity of single-cell data. As a forward-looking approach, scShiba will remain relevant and adaptable as the field evolves, bridging current technological limitations and future advancements. We are actively developing methods to better incorporate biological variability, aiming to further improve the tool’s functionality and applicability.

In conclusion, Shiba is a versatile well-around computational method for alternative splicing analysis, offering reliable detection of both annotated and unannotated DSEs. Its efficient runtime, reproducibility, consideration of junction read imbalance, and applicability to single-replicate and single-cell/nucleus data make it a valuable tool for researchers investigating transcriptome dynamics in diverse biological contexts. Shiba can make a significant contribution to the field of alternative splicing analysis and enhance our understanding of transcriptomic regulation.

## Author Contributions

Conceptualization: Naoto Kubota, Liang Chen, Sika Zheng. Formal analysis and software: Naoto Kubota. Funding acquisition: Liang Chen, Sika Zheng. Methodology: Liang Chen, Naoto Kubota. Project administration and resources: Sika Zheng. Supervision: Liang Chen, Sika Zheng. Writing-original draft: Naoto Kubota, Sika Zheng. Writing-review & editing: Naoto Kubota, Liang Chen, Sika Zheng.

## Acknowledgements

We thank all members of Sika Zheng’s laboratory for their valuable discussions. Manuscript writing was partially supported by ChatGPT to improve readability.

## Funding

This work was supported by National Institutes of Health [R01NS15276 to S.Z., R01GM137428 to L.C.]. N.K. was partially supported by the Uehara Memorial Foundation Postdoctoral Fellowship in 2022.

## Notes

### Competing Interest Statement

The authors have declared no competing interest.

### Summary of Updates

We simulated 104 different scenarios considering various combinations of read coverage, replicate numbers, and variations in replicate PSIs to compare Shiba with six other popular tools. We introduced a new metric, intra-to-inter ratio (IIR), to assess each tools specificity in analyzing real RNA-seq data. For RT-PCR validation, to avoid perceived bias from in-house experiments, we found two published RT-PCR results with numerical PSI values and previously used to validate results from popular methods like rMATS and MAJIQ. In both cases, Shiba outperformed the original methods. To further validate Shiba experimentally, we leverage AS-NMD transcripts, whose steady-state exon inclusion ratios and mRNA abundance are correlated, allowing us to (1) use changes in mRNA abundance upon NMD inhibition to corroborate changes in exon inclusion ratios, and (2) examine whether the identified changes in exon inclusion ratios upon NMD inhibition are associated with NMD annotation. We introduced Shiba+ to account for variability between replicates.

