## Supplemental_info for "Shiba: A versatile computational method for systematic identification of differential RNA splicing across platforms"

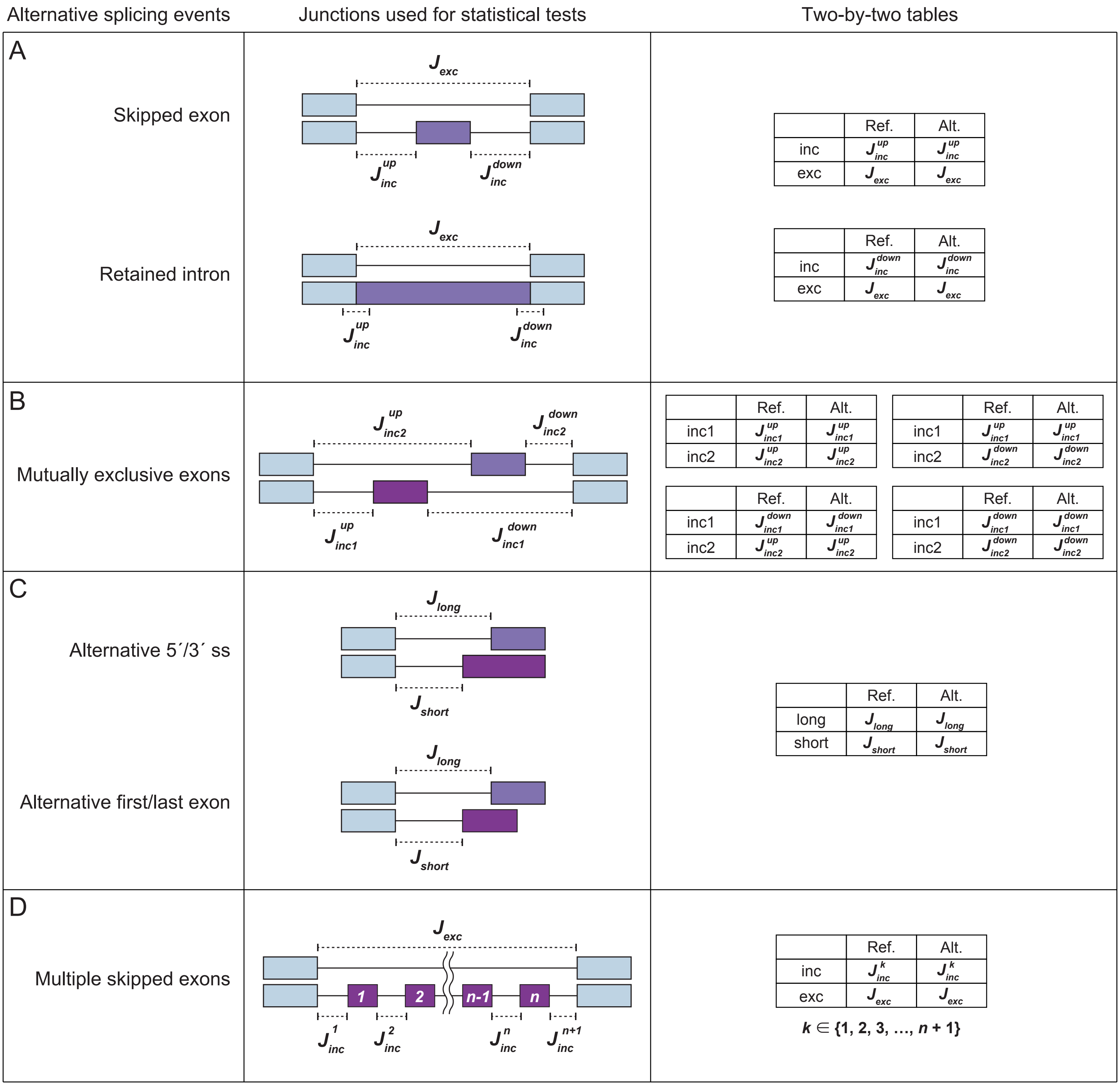


**Supplemental Figure 1. Schematic representation of exon-exon/exon-intron junctions used for statistical analysis in Shiba.** The two-by-two tables are used for Fisher’ s exact test. (A) Skipped exon and Retained intron, (B) Mutually exclusive exons, (C) Alternative 5´/3´ splice site and Alternative first/last exon, and (D) Multiple skipped exons.

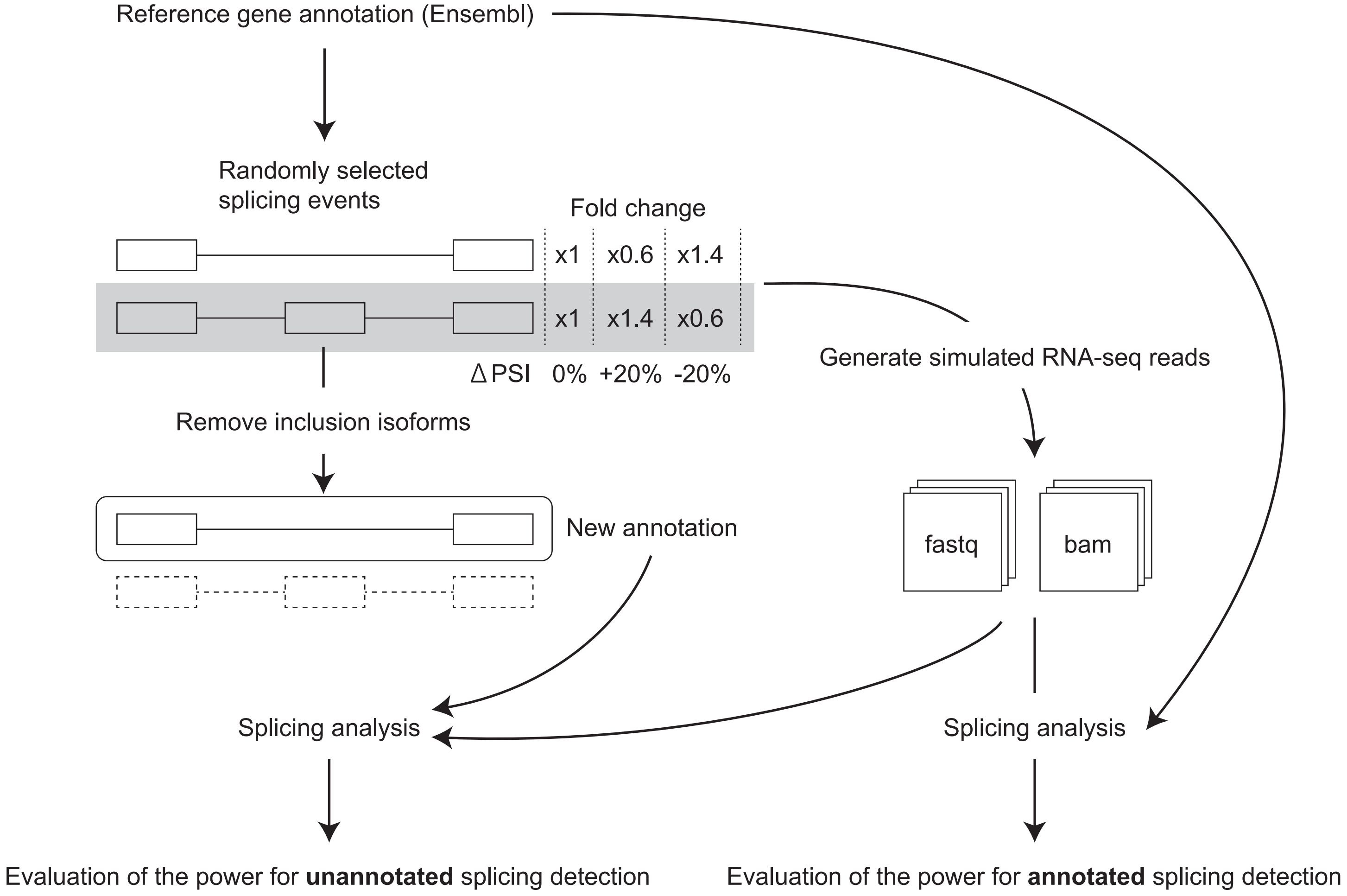


**Supplemental Figure 2. Generation of simulated RNA-seq data for detecting annotated and unannotated differential alternative splicing events.** We randomly selected mouse protein-coding genes that have alternative splicing events (SE, FIVE, THREE, MXE, RI, MSE, AFE, and ALE) and divided them into three groups: (i) Unaltered genes, (ii) Genes with 20% PSI increase, and (iii) Genes with 20% PSI decrease in the perturbed samples. For the performance evaluation of detecting unannotated alternative splicing events, we customized the reference gene annotation file by removing records of inclusion isoforms.

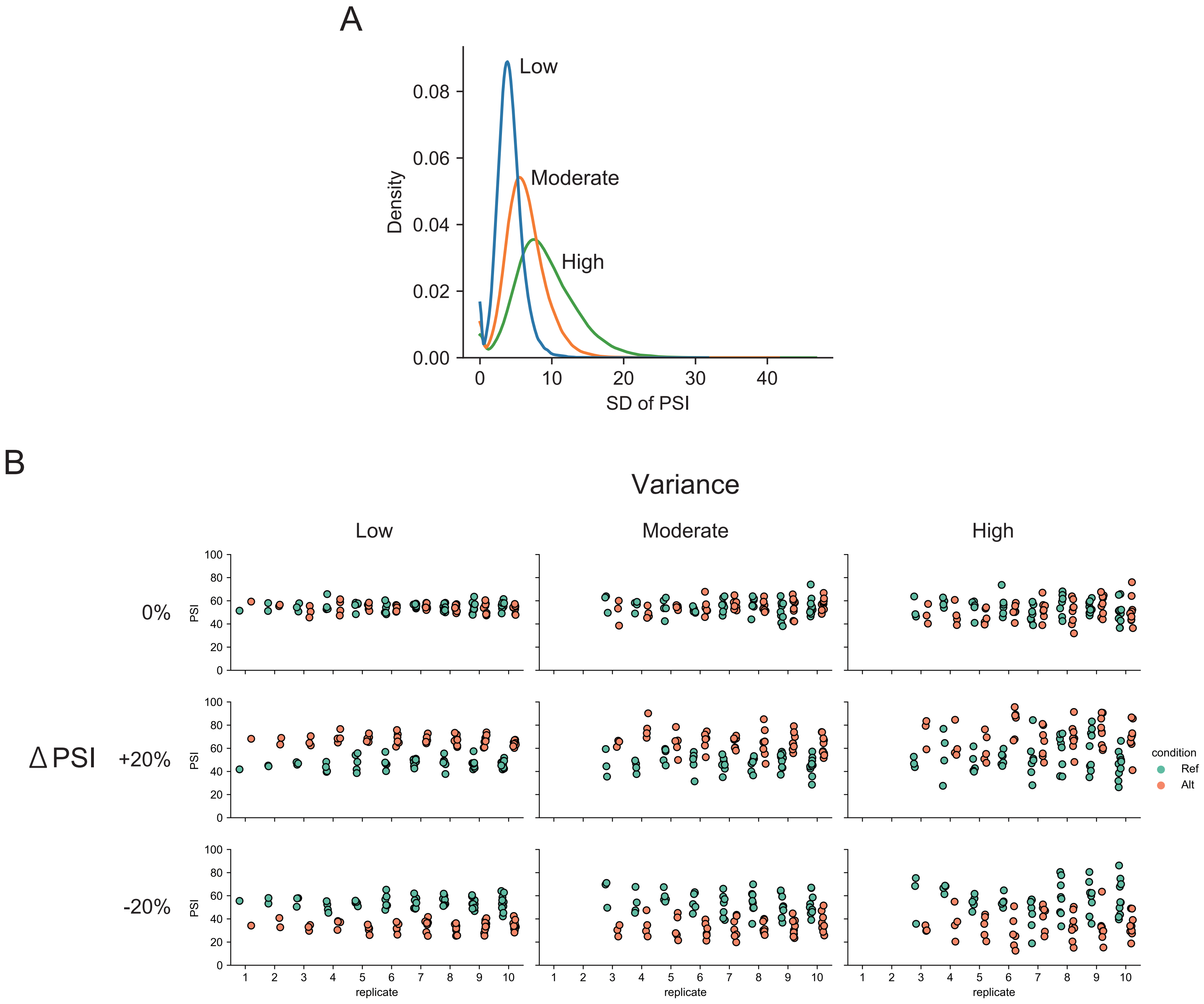


**Supplemental Figure 3. PSI in simulated RNA-seq data used performance evaluation.** (A) Distribution of standard deviation of percent spliced in (PSI) for individual samples in low, moderate, and high variance conditions. (B) PSI of example events that exhibit unaltered or altered splicing (20% increase or decrease) in individual samples with low, moderate, and high variance.

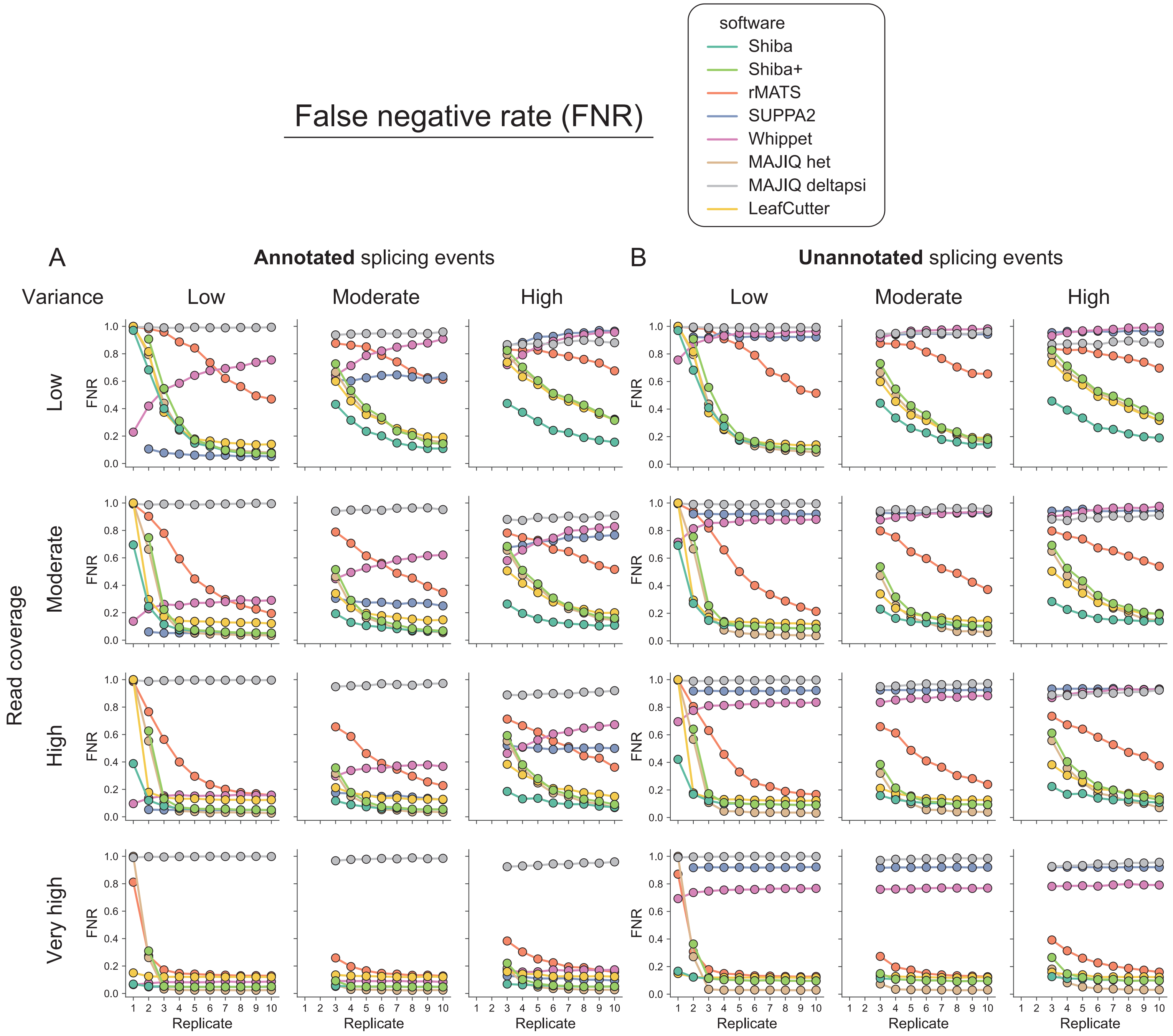


**Supplemental Figure 4. False negative rate in simulated RNA-seq data for five classical splicing events.** (A) False negative rate (FNR) for annotated splicing event detection by eight computational tools/modules (Shiba, Shiba+, rMATS, SUPPA2, Whippet, MAJIQ het, MAJIQ deltapsi, and LeafCutter) in different combinations of sample variations (Low, Moderate, and High), read coverages (Low, Moderate, High, and Very high) and biological replicate numbers (1 ~ 10). (B) The corresponding metric for unannotated splicing event detection.

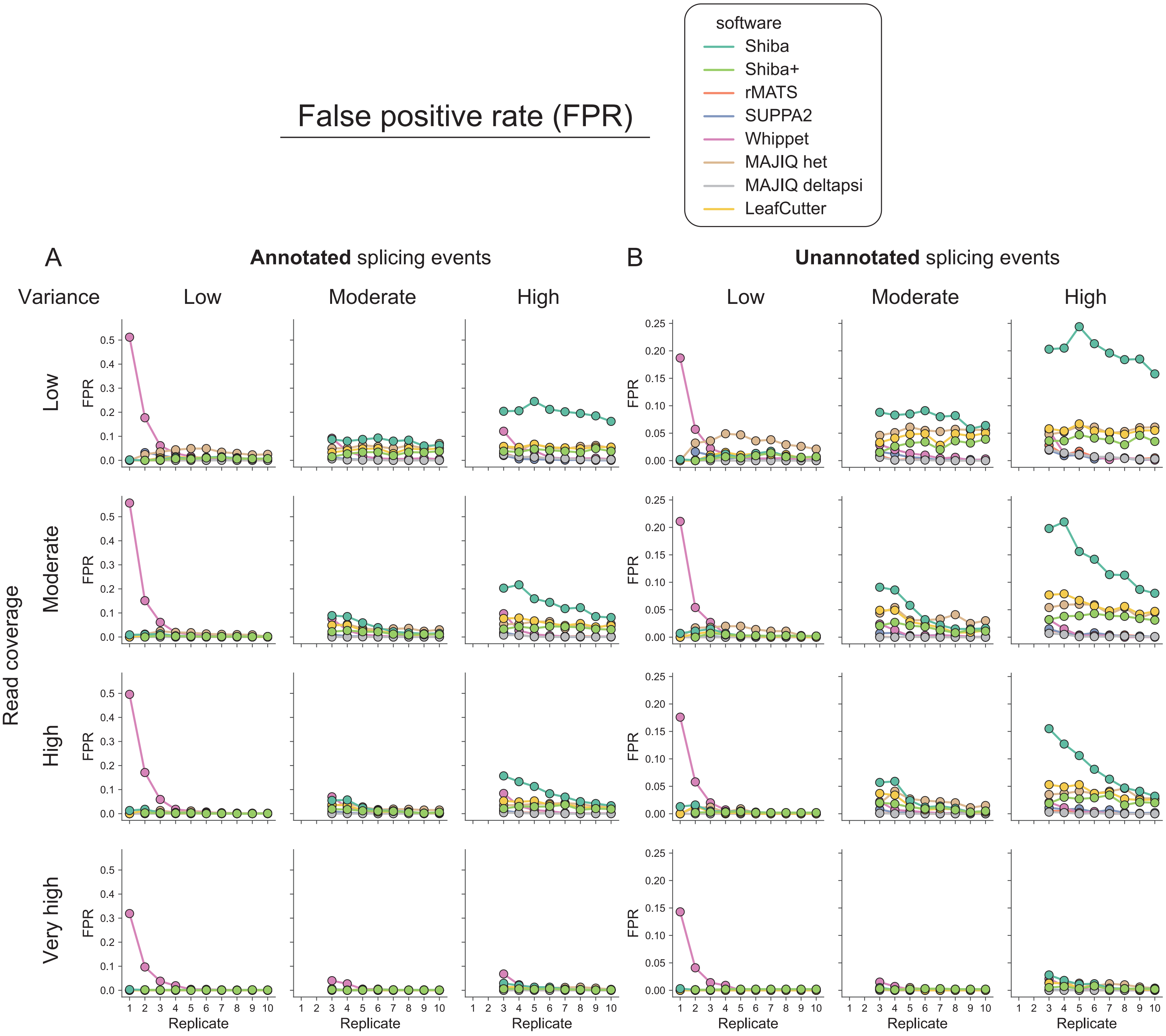


**Supplemental Figure 5. False positive rate in simulated RNA-seq data for five classical splicing events.** (A) False positive rate (FPR) for annotated splicing event detection by eight computational tools/modules (Shiba, Shiba+, rMATS, SUPPA2, Whippet, MAJIQ het, MAJIQ deltapsi, and LeafCutter) in different combinations of sample variations (Low, Moderate, and High), read coverages (Low, Moderate, High, and Very high) and biological replicate numbers (1 ~ 10). (B) The corresponding metric for unannotated splicing event detection.

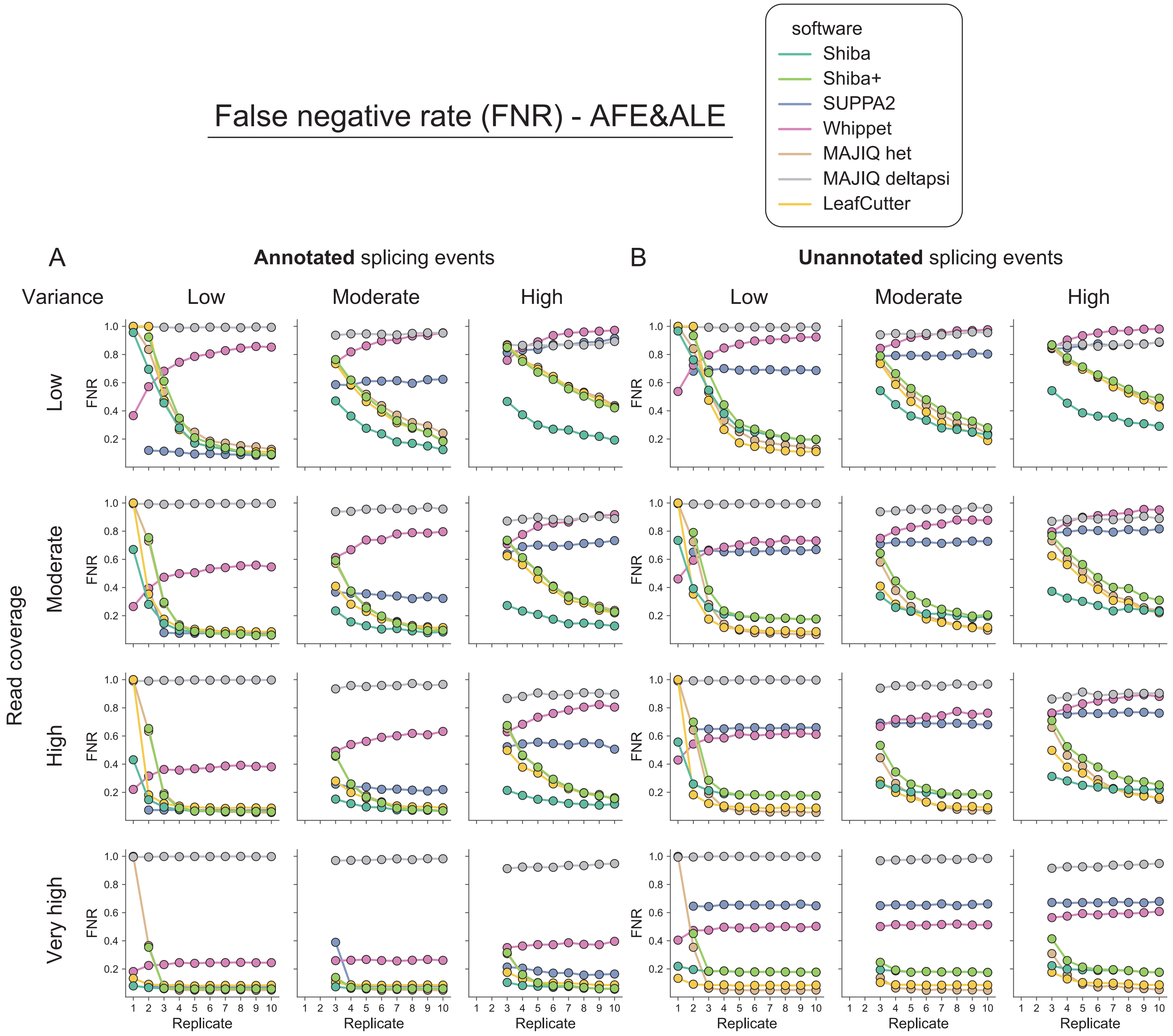


**Supplemental Figure 6. False negative rate in simulated RNA-seq data for alternative first and last exons.** (A) False negative rate (FNR) for annotated alternative first and last exon detection by seven computational tools/modules (Shiba, Shiba+, SUPPA2, Whippet, MAJIQ het, MAJIQ deltapsi, and LeafCutter) in different combinations of sample variations (Low, Moderate, and High), read coverages (Low, Moderate, High, and Very high) and biological replicate numbers (1 ~ 10). (B) The corresponding metric for unannotated splicing event detection.

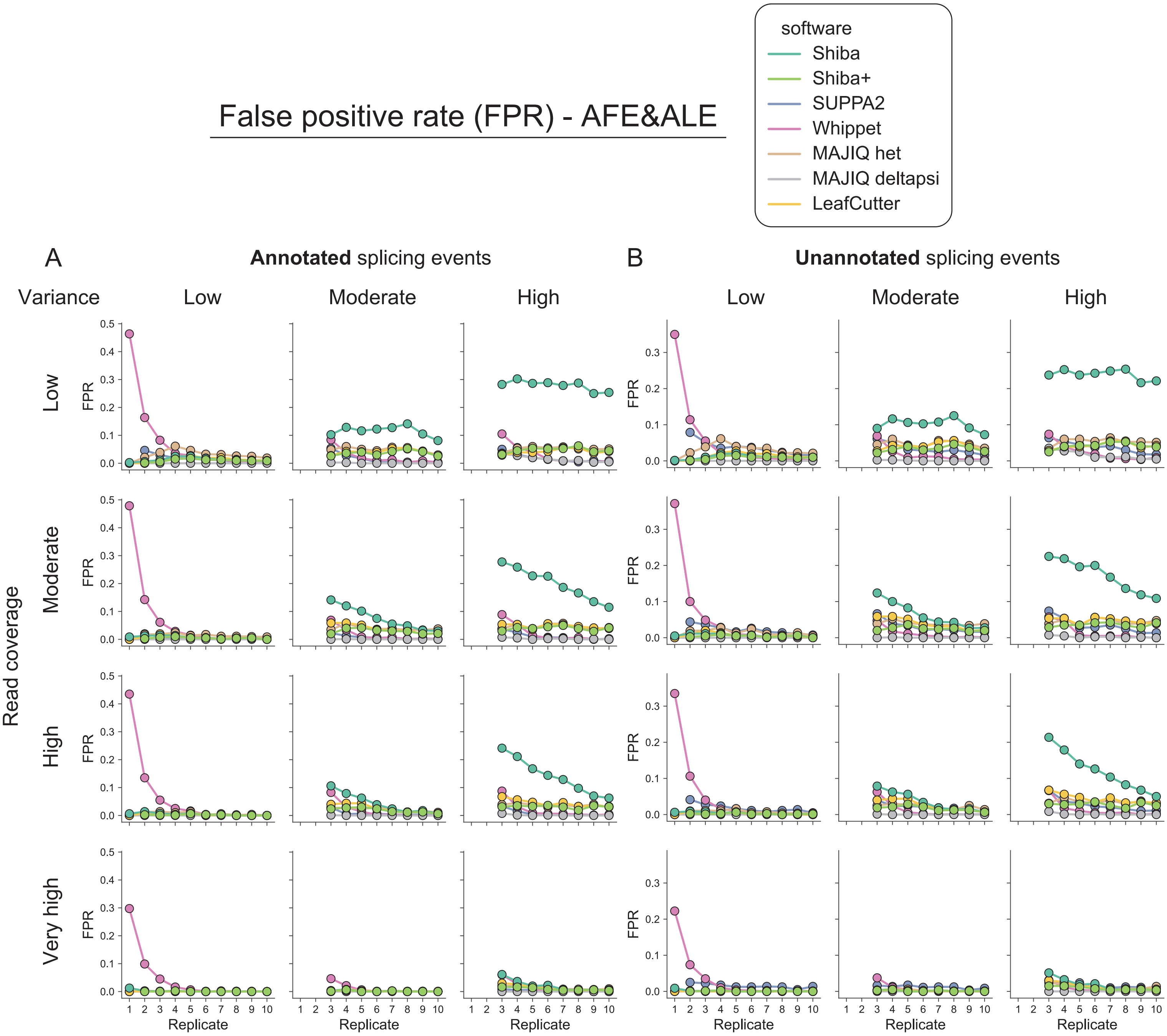


**Supplemental Figure 7. False positive rate in simulated RNA-seq data for alternative first and last exons.** (A) False positive rate (FPR) for annotated alternative first and last exon detection by seven computational tools/modules (Shiba, Shiba+, SUPPA2, Whippet, MAJIQ het, MAJIQ deltapsi, and LeafCutter) in different combinations of sample variations (Low, Moderate, and High), read coverages (Low, Moderate, High, and Very high) and biological replicate numbers (1 ~ 10). (B) The corresponding metric for unannotated splicing event detection.

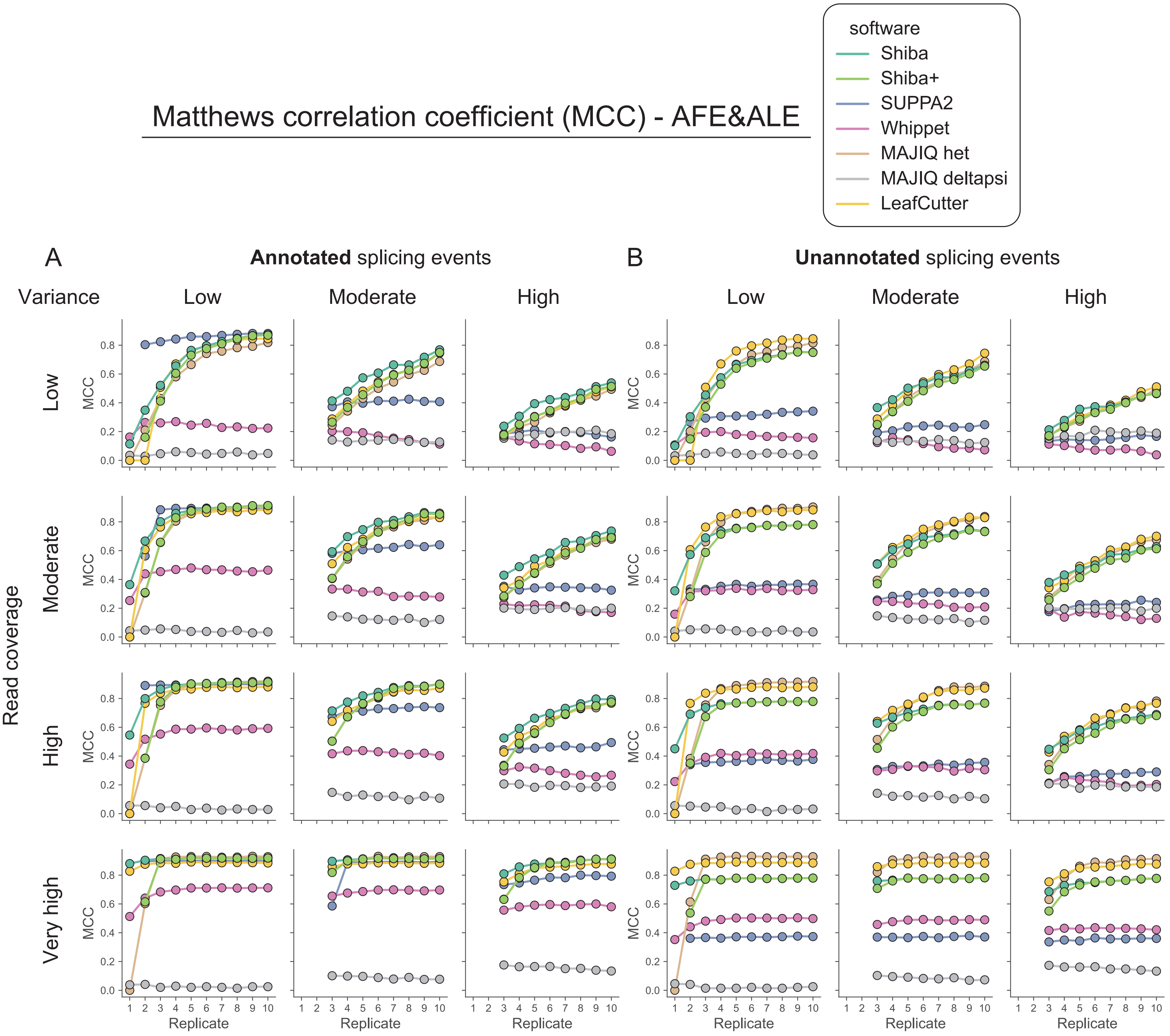


**Supplemental Figure 8: Matthews correlation coefficient in simulated RNA-seq data for alternative first and last exons.** (A) Matthews correlation coefficient (MCC) for annotated alternative first and last exon detection by seven computational tools/modules (Shiba, Shiba+, SUPPA2, Whippet, MAJIQ het, MAJIQ deltapsi, and LeafCutter) in different combinations of sample variations (Low, Moderate, and High), read coverages (Low, Moderate, High, and Very high) and biological replicate numbers (1 ~ 10). (B) The corresponding metric for unannotated splicing event detection.

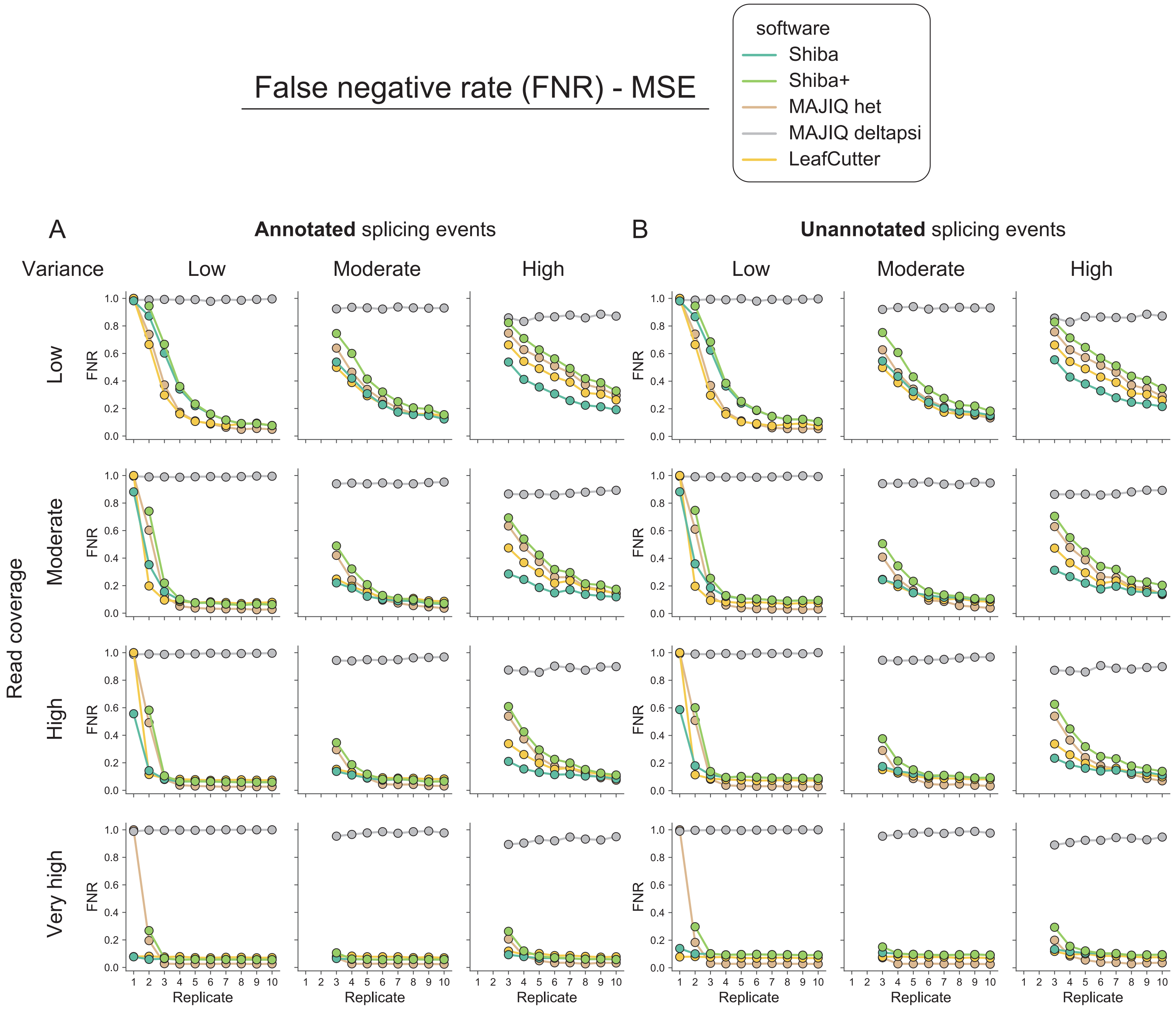


**Supplemental Figure 9. False negative rate in simulated RNA-seq data for multiple skipped exons.** (A) False negative rate (FNR) for annotated multiple skipped exon detection by five computational tools/modules (Shiba, Shiba+, MAJIQ het, MAJIQ deltapsi, and LeafCutter) in different combinations of sample variations (Low, Moderate, and High), read coverages (Low, Moderate, High, and Very high) and biological replicate numbers (1 ~ 10). (B) The corresponding metric for unannotated splicing event detection.

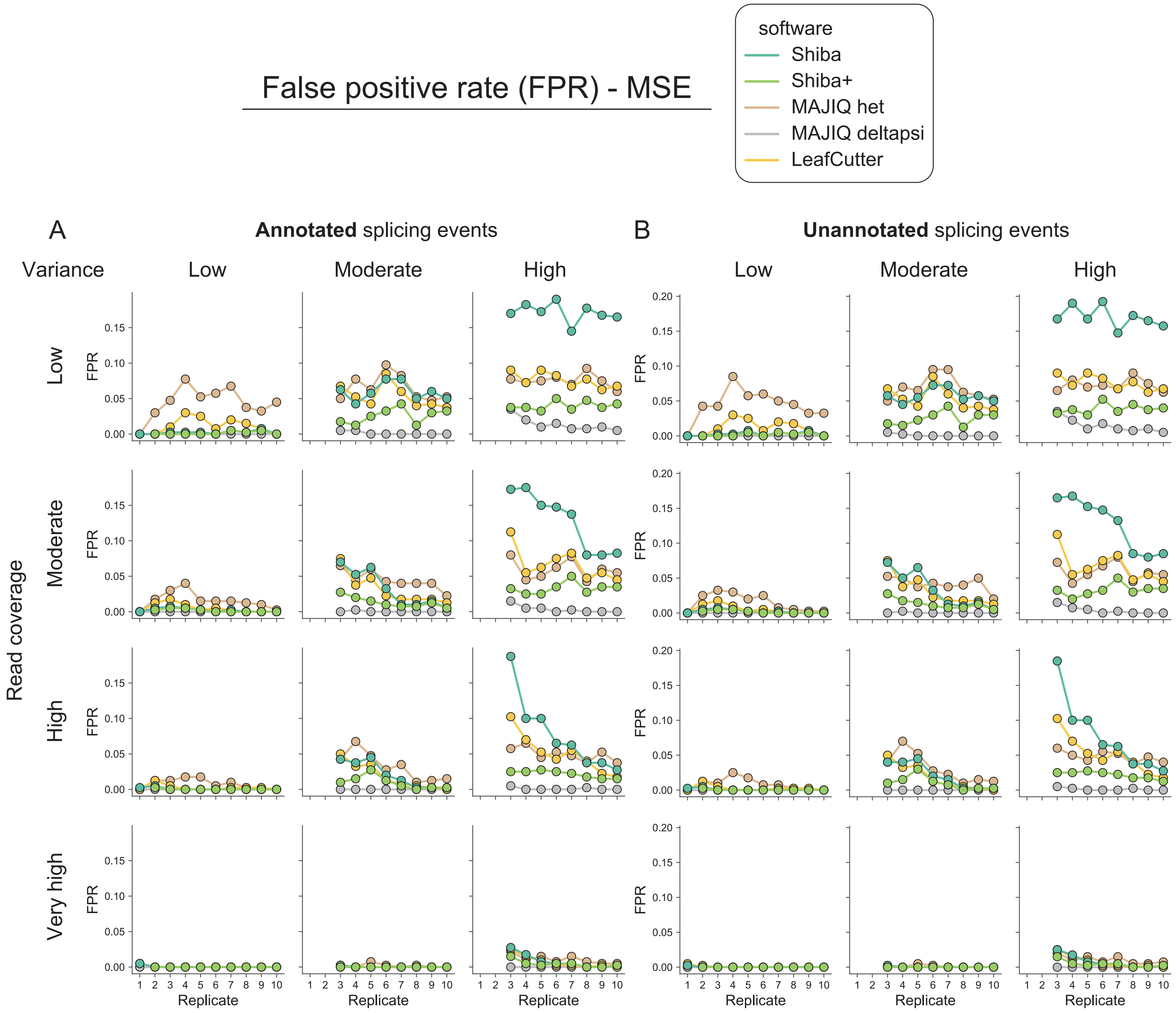


**Supplemental Figure 10. False positive rate in simulated RNA-seq data for multiple skipped exons.** (A) False positive rate (FPR) for annotated multiple skipped exon detection by five computational tools/modules (Shiba, Shiba+, MAJIQ het, MAJIQ deltapsi, and LeafCutter) in different combinations of sample variations (Low, Moderate, and High), read coverages (Low, Moderate, High, and Very high) and biological replicate numbers (1 ~ 10). (B) The corresponding metric for unannotated splicing event detection.

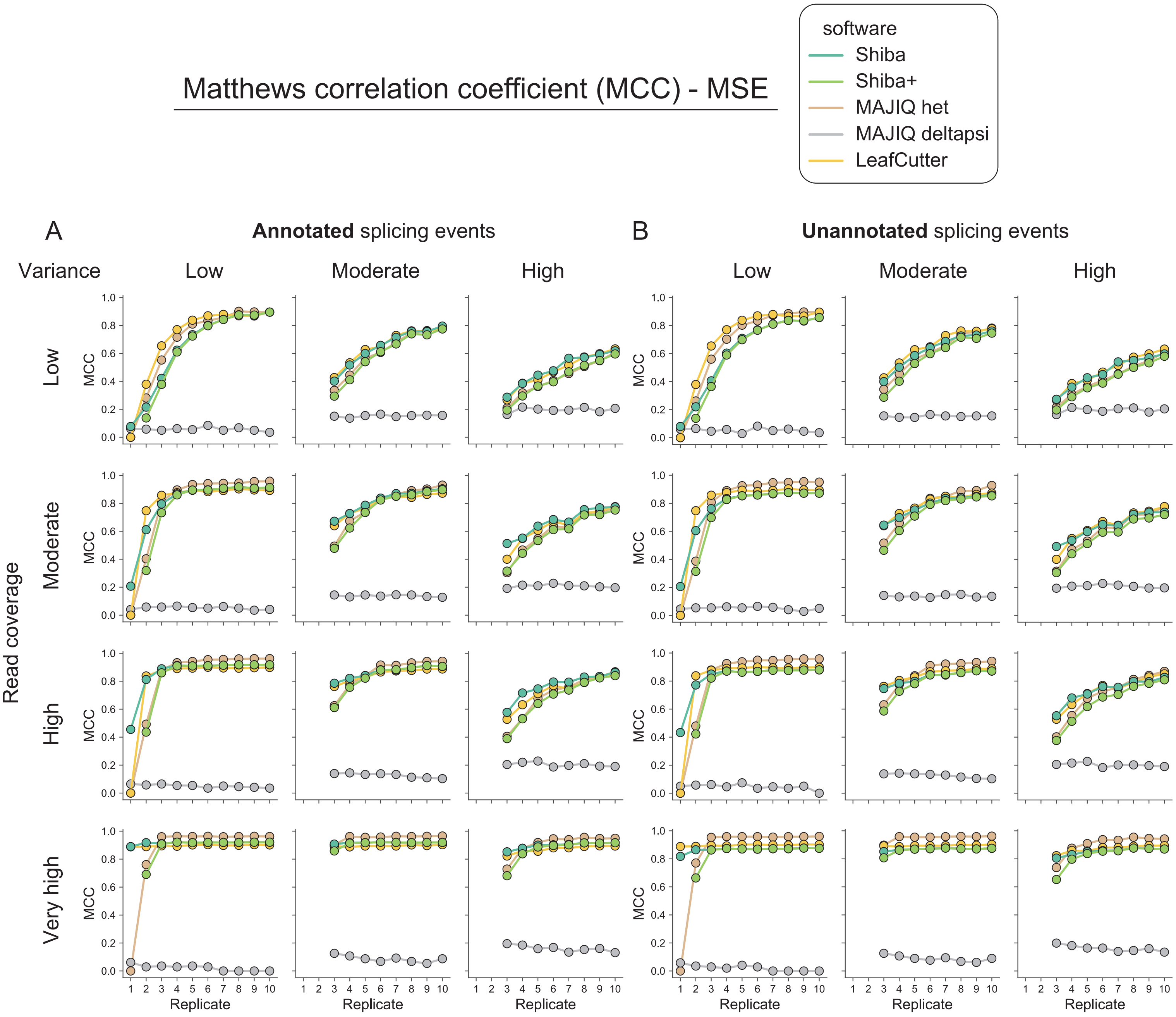


**Supplemental Figure 11: Matthews correlation coefficient in simulated RNA-seq data for multiple skipped exons.** (A) Matthews correlation coefficient (MCC) for annotated multiple skipped exon detection by five computational tools/modules (Shiba, Shiba+, MAJIQ het, MAJIQ deltapsi, and LeafCutter) in different combinations of sample variations (Low, Moderate, and High), read coverages (Low, Moderate, High, and Very high) and biological replicate numbers (1 ~ 10). (B) The corresponding metric for unannotated splicing event detection.

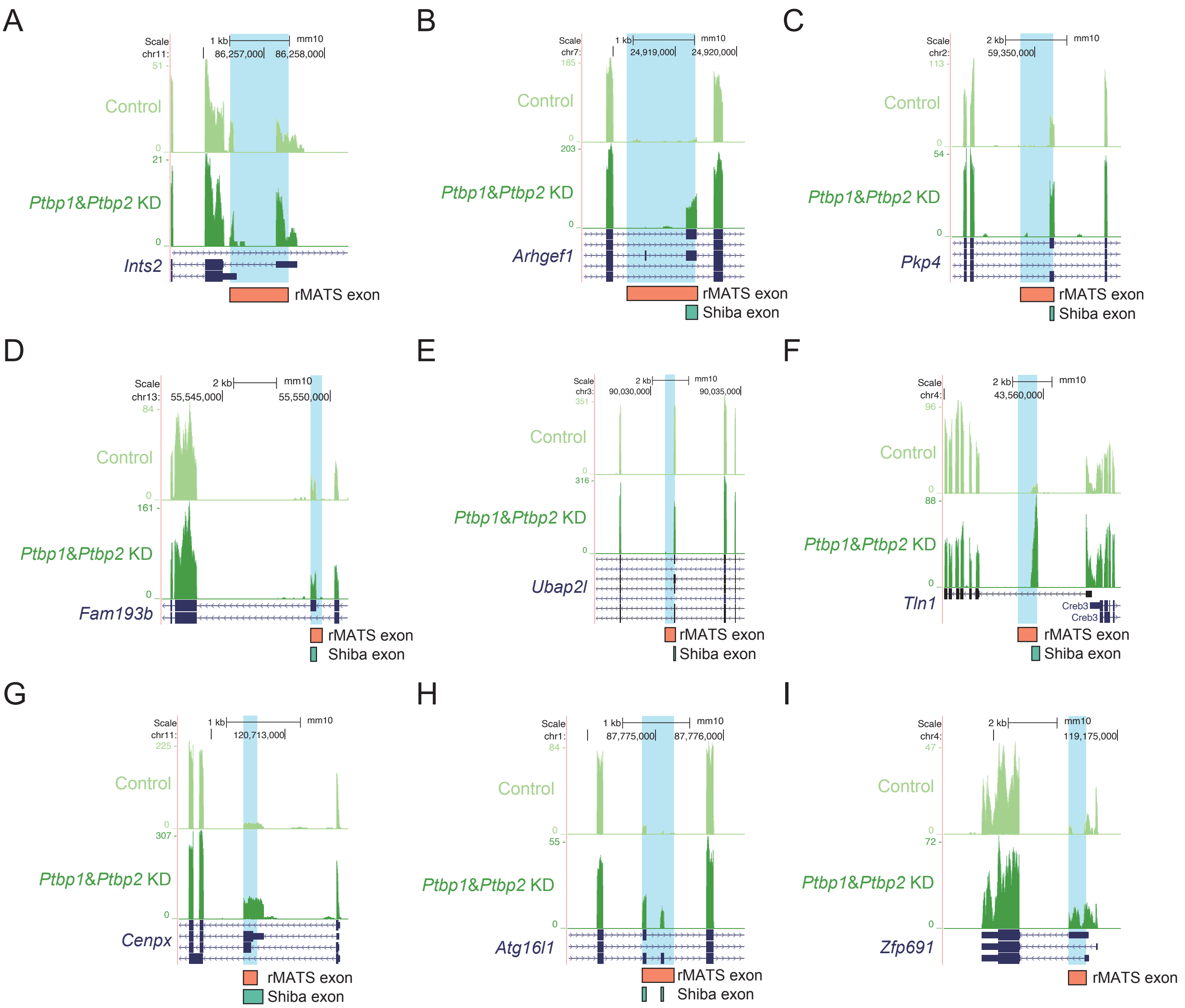


**Supplemental Figure 12. Examples of rMATS-identified events with potential errors in exon definition.** Detected alternative exons in (A) Ints2, (B) Arhgef1, (C) Pkp4, (D) Fam193b, (E) Ubap2l, (F) Tln1, (G) Cenpx, (H) Atg16l1 and (I) Zfp691 are shown with read coverage tracks of RNA-seq samples from the cytoplasmic fraction of mESC with siRNA-mediated double knockdown of Ptbp1 and Ptbp2 [Yeom et al., 2021]. Exons identified by rMATS and Shiba are shown in the orange and green colored boxes, respectively.

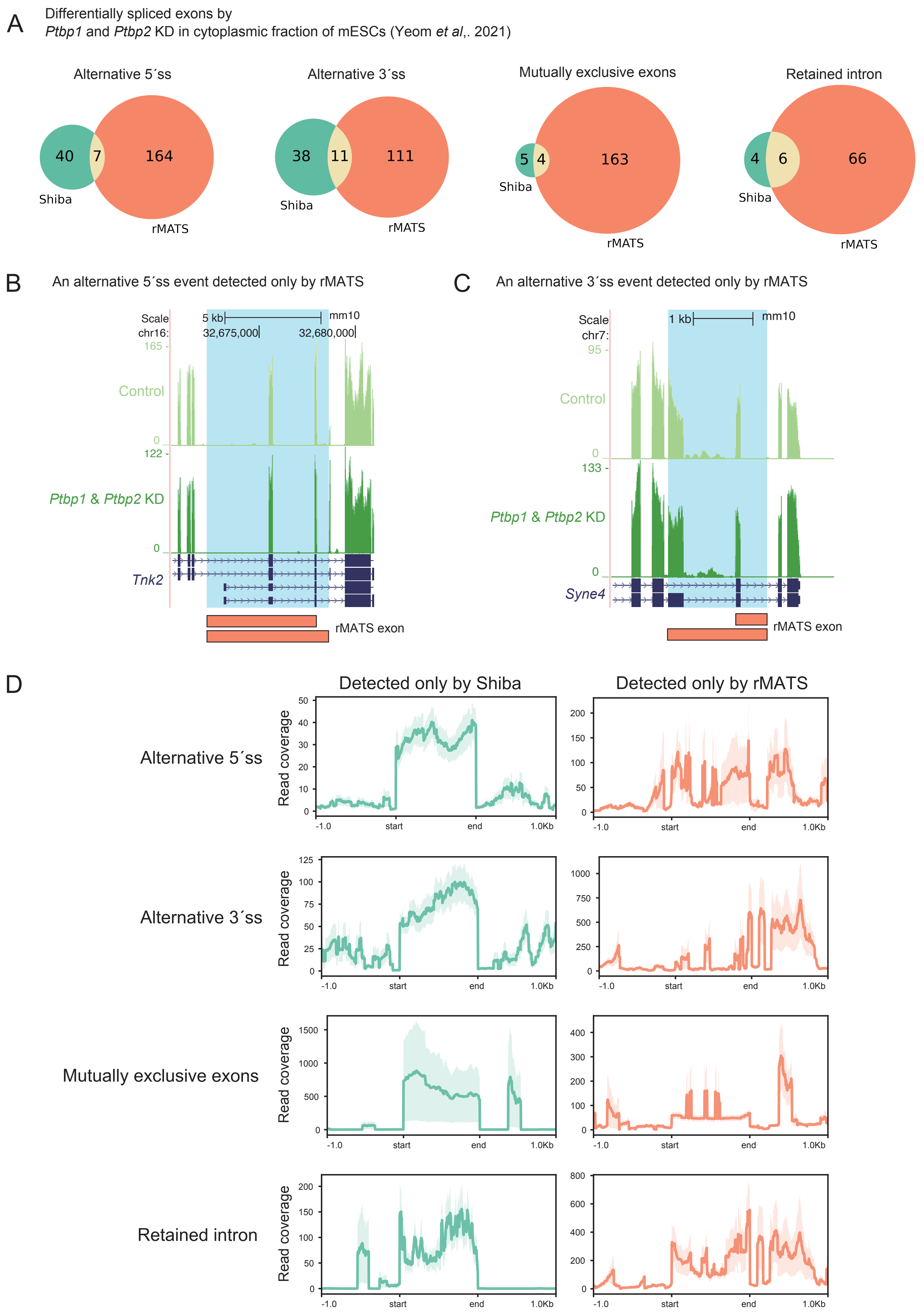

A
Differentially spliced exons by
Ptbp1 and Ptbp2 KD in cytoplasmic fraction of mESCs (Yeom et al,. 2021)
Alternative 5´ss
40
7
164
Shiba
rMATS
Alternative 3´ss
38
11
111
Shiba
rMATS
Mutually exclusive exons
5
4
163
Shiba
rMATS
Retained intron
4
6
66
Shiba
rMATS
B
An alternative 5´ss event detected only by rMATS
Scale
5 kb
mm10
chr16:
32,675,000
32,680,000
Control
Ptbp1 & Ptbp2 KD
Tnk2
rMATS exon
C
An alternative 3´ss event detected only by rMATS
Scale
1 kb
mm10
chr7:
Control
Ptbp1 & Ptbp2 KD
Syne4
rMATS exon
D
Detected only by Shiba
Detected only by rMATS
Alternative 5´ss
Alternative 3´ss
Mutually exclusive exons
Retained intron
Read coverage
-1.0
start
end
1.0Kb

**Supplemental Figure 13. Comparison of differentially spliced events detected by Shiba vs. rMATS.** (A) Comparison of alternative five prime splice site (FIVE), alternative three prime splice site (THREE), mutually exclusive exons (MXE), and retained intron (RI) events detected by Shiba and rMATS in the cytoplasmic fraction of mESC with siRNA-mediated double knockdown of Ptbp1 and Ptbp2 [Yeom et al., 2021]. (B) A specific FIVE event in the Tnk2 gene, exclusively identified by rMATS, shows potential mis-annotation illustrated by discrepancy from the RNA-seq browser tracks. The orange boxes represent the longer and shorter exons defined by rMATS. (C) A specific THREE event in the Syne4 gene exclusively identified by rMATS. (D) Aggregation plots of read coverage on the meta-exon for FIVE, THREE, MXE, and RI, demonstrating even distribution for Shiba-identified events and uneven distribution for rMATS-identified events.

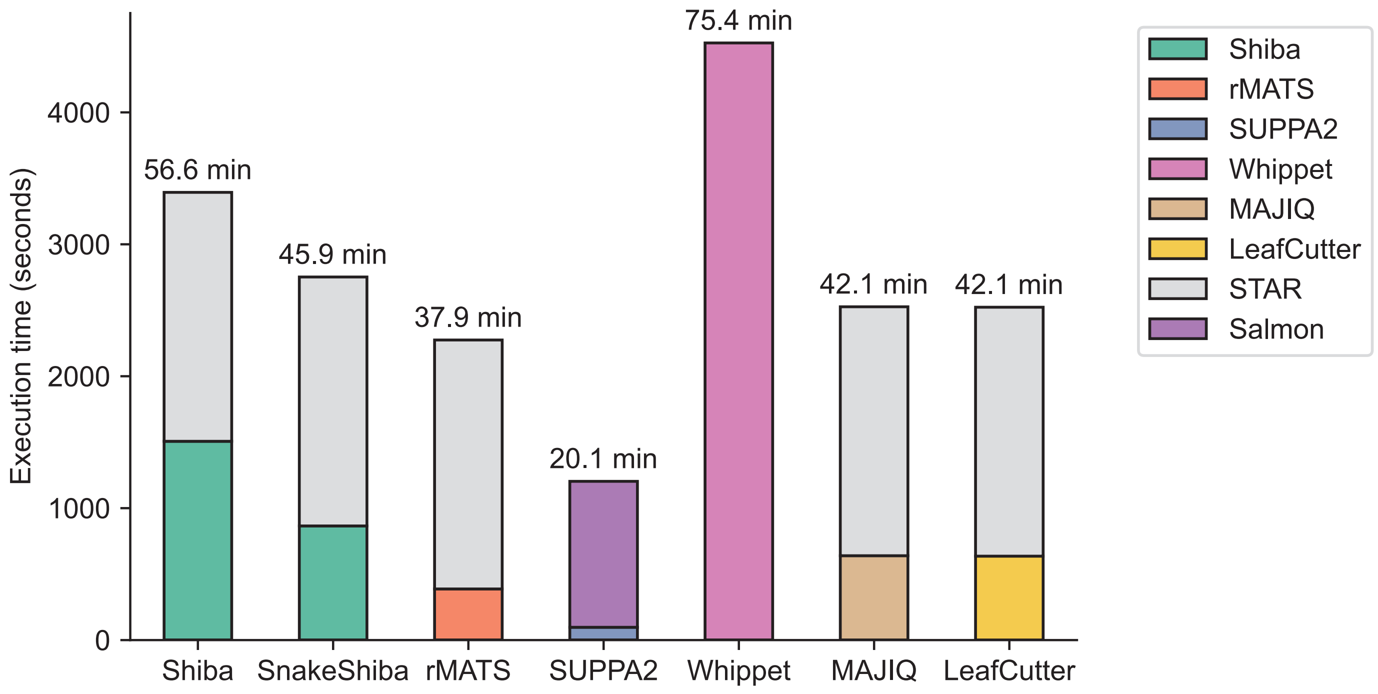


**Supplemental Figure 14. Runtime comparison of computational tools for differential splicing analysis using real RNA-seq data.** The runtime assessment utilized a dataset from Yeom et al. [Yeom *et al*., 2021], featuring control and perturbed samples with three biological replicates, providing a representative dataset. Sixteen CPUs were allocated for Shiba, SnakeShiba, rMATS, MAJIQ, LeafCutter, STAR, and Salmon, while SUPPA2 and Whippet, lacking multi-core support, were executed on a single CPU.

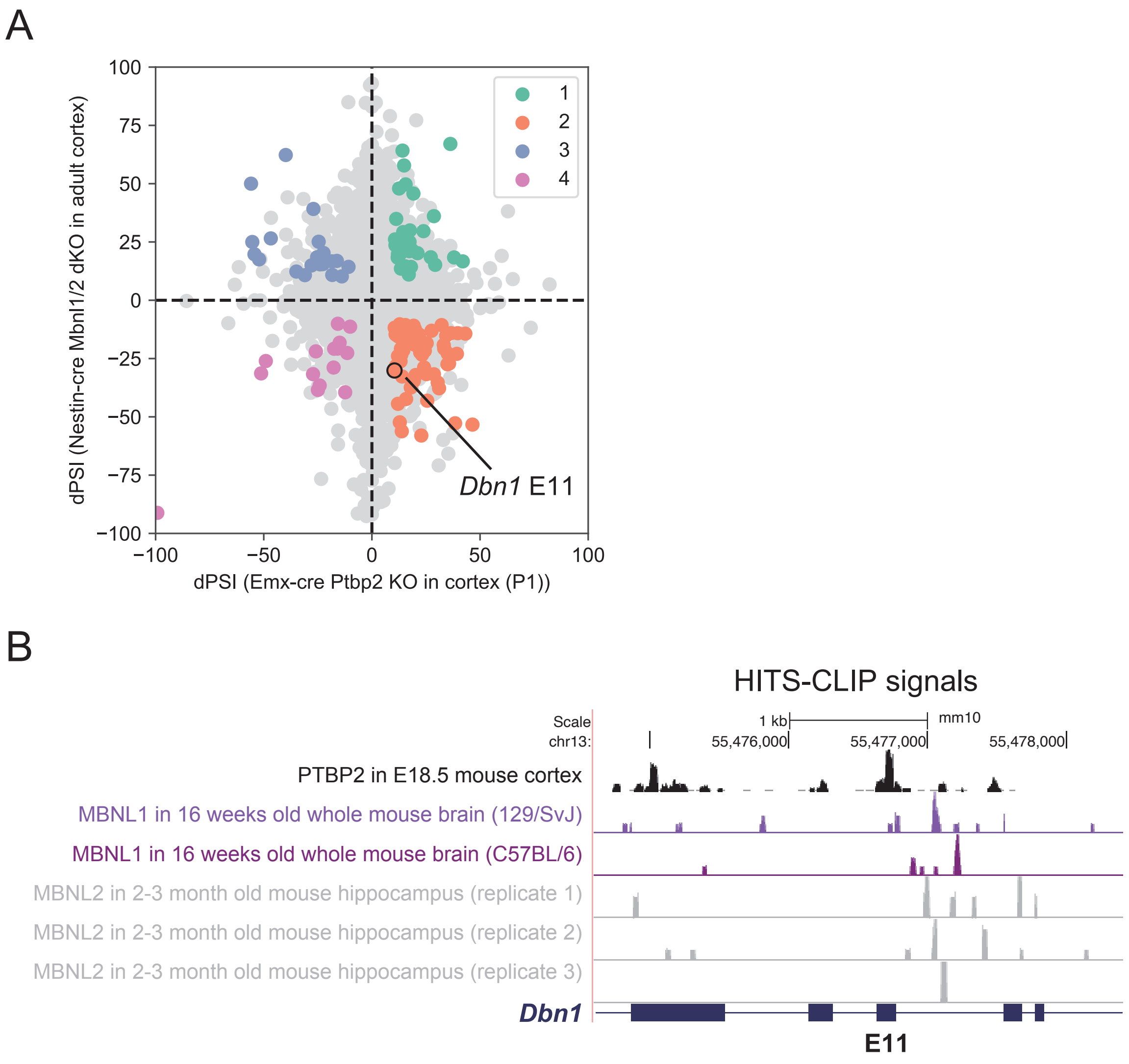


**Supplemental Figure 15. Alternative splicing events under combinatorial Ptbp2 and Mbnl1/2 regulation.** (A) Scatter plot of PSI changes in Emx1-cre Ptbp2 KO (x-axis) and Nestin-cre Mbnl1/2 dKO (y-axis). Colored dots represent events claimed as differentially spliced events (DSEs) by Shiba ($q < 0.05$ and $|\Delta PSI| > 10$) in both conditions. Group 1 (green): Up-regulated in both, Group 2 (orange): Up-regulated in Ptbp2 KO and down-regulated in Mbnl1/2 dKO, Group 3 (blue): Down-regulated in Ptbp2 KO and up-regulated in Mbnl1/2 dKO, Group 4 (pink): Down-regulated in both. Dbn1 E11 is marked in group 2. (B) Genome browser track of HITS-CLIP signals around the genomic region of Dbn1 E11.

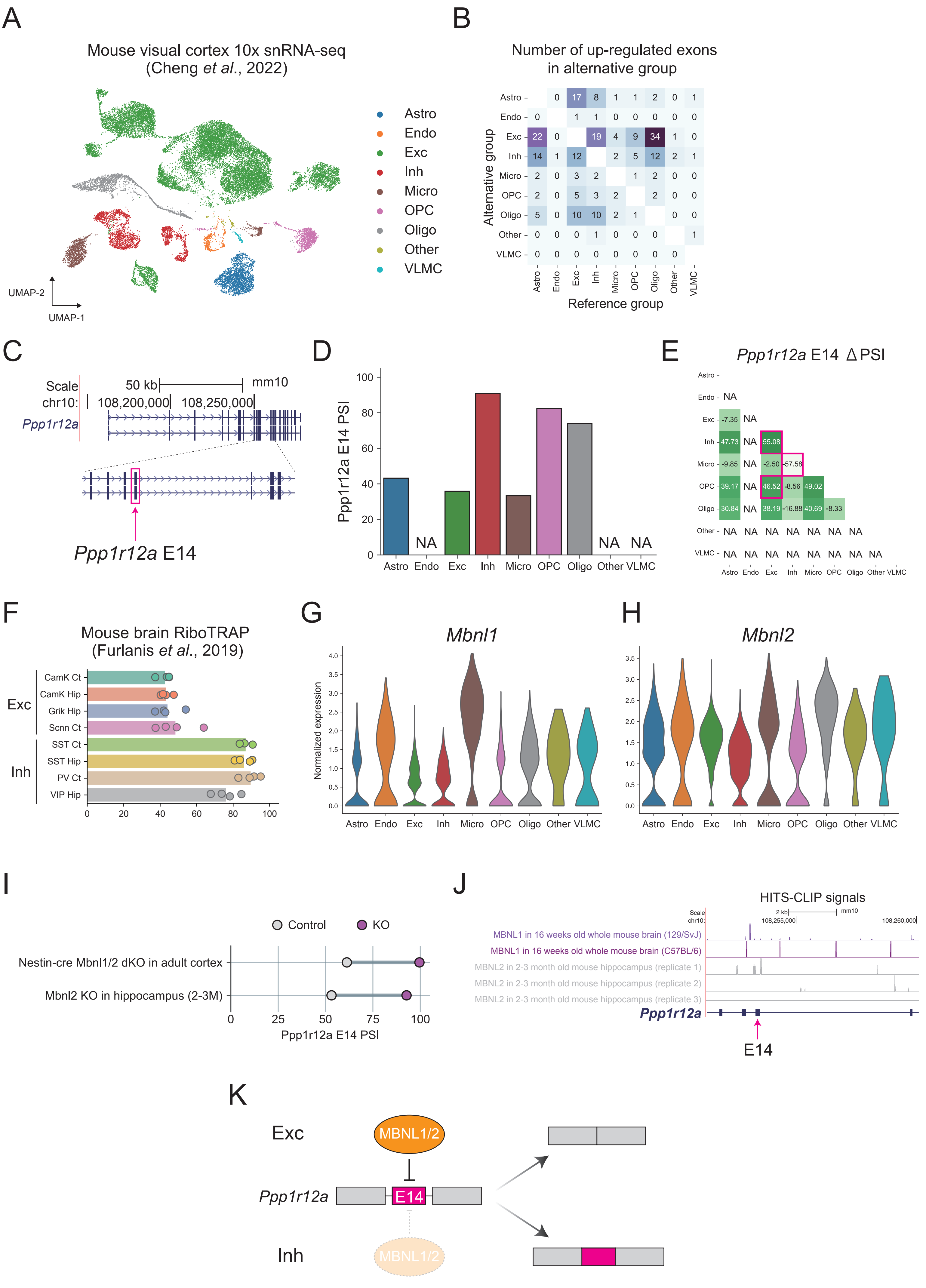
A
Mouse visual cortex 10x snRNA-seq (Cheng et al., 2022)
Astro
Endo
Exc
Inh
Micro
OPC
Oligo
Other
VLMC
UMAP-2
UMAP-1
B
Number of up-regulated exons in alternative group
Alternative group
Reference group
C
Scale
50 kb
mm10
chr10:
108,200,000
108,250,000
Ppp1r12a
Ppp1r12a E14
D
Ppp1r12a E14 PSI
NA
E
Ppp1r12a E14 ΔPSI
F
Mouse brain RiboTRAP (Furlanis et al., 2019)
CamK Ct
CamK Hip
Grik Hip
Scnn Ct
SST Ct
SST Hip
PV Ct
VIP Hip
G
Mbnl1
Normalized expression
H
Mbnl2
I
Control
KO
Nestin-cre Mbnl1/2 dKO in adult cortex
Mbnl2 KO in hippocampus (2-3M)
Ppp1r12a E14 PSI
J
HITS-CLIP signals
MBNL1 in 16 weeks old whole mouse brain (129/SvJ)
MBNL1 in 16 weeks old whole mouse brain (C57BL/6)
MBNL2 in 2-3 month old mouse hippocampus (replicate 1)
MBNL2 in 2-3 month old mouse hippocampus (replicate 2)
MBNL2 in 2-3 month old mouse hippocampus (replicate 3)
E14
K
Exc
MBNL1/2
E14
Inh

**Supplemental Figure 16. scShiba identifies cell-type-specific alternative splicing events using 10x single-nucleus RNA-seq data.** (A) 10x single-nucleus (sn) RNA-seq from mouse visual cortex [Cheng et al., 2022], plotted by the first two dimensions of UMAP. Astro: Astrocyte, Endo: Endothelial cell, Exc: Excitatory neuron, Inh: Inhibitory neuron, Micro: Microglia, OPC: Oligodendrocyte progenitor cell, Oligo: Oligodendrocyte, Other; Other cell types, VLMC: Vascular leptomeningeal cell. (B) A heatmap of number of differentially spliced events (DSEs) detected in scShiba. The number in each box represents up-regulated DSEs ($q < 0.05$, $\Delta PSI > 10$) in the alternative group compared to the reference group. (C) Genome browser track of Ppp1r12a gene locus. (D) PSI values of Ppp1r12a E14 in each cell type from the snRNA-seq data. (E) ΔPSI values of Ppp1r12a E14 in all comparisons between cell types. Magenta boxes represent statistical significance ($q < 0.05$, $|\Delta PSI| > 10$). (F) PSI values of Ppp1r12a E14 in eight neuronal subtypes of excitatory and inhibitory neurons from RiboTRAP data [Furlanis et al., 2019]. Camk Ct: CamK2+ neurons from cortex, Camk Hip: CamK2+ neurons from hippocampus, Grik Hip: Grik4+ neurons from hippocampus, Scnn Ct: Scnn1a+ neurons from cortex, SST Ct: SST+ neurons from cortex, SST Hip: SST+ neurons from hippocampus, PV Ct: PV+ neurons from cortex, VIP Hip: VIP+ neurons from hippocampus. (G-H) Expression levels of Mbnl1 (G) and Mbnl2 (H) for each cell type in the snRNA-seq data. (I) PSI values of Ppp1r12a E14 in control and KO samples from Nestin-cre Mbnl1 and Mbnl2 double KO mouse adult cortex [Vanhentenryck et al., 2018] and Mbnl2 KO mouse adult hippocampus (2–3 months old) [Charizanis et al., 2012]. (J) Genome browser track of HITS-CLIP signals around the genomic region of Ppp1r12a E14. (K) A model of alternative splicing program of Ppp1r12a E14 in excitatory and inhibitory neurons.

**Strengths of Shiba and Shiba+**

| Comparison matrix | Reference figure |
|---|---|
| Consistent and high reproducibility ratio | Figure 3A-B |
| Low Intra-to-inter ratio from real data (i.e., high specificity) | Figure 3C-D |
| High correlation with RT-PCR in ΔPSI quantification | Figure 3E-F |
| High MCC from simulated RNA-seq data | Figure 2; Supplementary Figure S8, S11 |
| Low FNR from simulated RNA-seq data in different scenarios | Supplementary Figure S4, S6, S9 |

**Features of the methods**

| | Reference figures | Shiba | Shiba+ | rMATS | SUPPA2 | Whippet | MAJIQ het | MAJIQ deltapsi | LeafCutter |
|---|---|---|---|---|---|---|---|---|---|
| Consideration of junction read imbalance | Figure 4 | Yes | Yes | No | No | No | NA | NA | NA |
| Detection of alternative first/last exons | Supplementary Figure S6-8 | Yes | Yes | No | Yes | Yes | Yes | Yes | Yes |
| Detection of multiple skipped exons | Supplementary Figure S9-11 | Yes | Yes | No | No | No | Yes | Yes | Yes |
| Detection of unannotated splicing events | Figure 2B; Supplementary Figure S4B, S5B, S6B, S7B, S8B, S9B, S10B, S11B | Yes | Yes | Yes | No | No | Yes | Yes | Yes |
| Consideration of exon body coverage | Figure 4I; Supplementary Figure S13D | Yes | Yes | No | Yes | No | No | No | No |
| Effectiveness in analyzing $n=1$ RNA-seq data | Figure 2, Figure 5 | High | does not support | Ineffective | does not support | Ineffective | does not support | Low | High |
